# Study of Principles Governing Epithelial Cell Clustering and Collective Motion *In Vitro*

**DOI:** 10.64898/2026.07.29.741528

**Authors:** Jia Gou, Mykhailo Potomkin, Jonard P. Ingal, Sergei Butenko, Wendy F. Liu, Maksim V. Plikus, Mark Alber

**Affiliations:** Department of Mathematics, University of California, Riverside, CA, United States; Interdisciplinary Center for Data-Driven Modeling in Biology, University of California, Riverside, CA, United State; Department of Developmental and Cell Biology, University of California, Irvine, CA, United States; Department of Biomedical Engineering, University of California, Irvine, Irvine, California, USA; A*STAR Skin Research Labs (A*SRL), Singapore, Republic of Singapore

## Abstract

In epithelial wounds, physically enlarged cells emerge along injury edges where they interact with regular-size cells during collective migration to repair tissue continuity. Although such large cells are often interpreted as migration leaders, recent observations suggest that regular-size cells can reciprocally influence them and that mixed-size cell clusters engage in distinct rotational motion before merging into confluent sheets. How cell size, polarity and adhesive coupling jointly control distinct collective migration modes remains unclear. Because many features of collective epithelial migration, including large and regular-size cell phenotypes, are conserved between *in vivo* and *in vitro* systems, here we developed a multi-scale computational model of interacting epithelial cells in two-dimensional culture with dynamic cell-substrate adhesion, cell-cell interactions, protrusion-based polarity, and contact-induced myosin redistribution as polarity regulator. We also explicitly modeled two functionally distinct cell states, featuring regular and large sizes respectively. Simulations showed that symmetric and asymmetric myosin redistribution at cell-cell junctions can produce stable cell contacts and persistent rotational motions by cell doublets. Intriguingly, in mixed-size cell clusters, straight translation can arise both from leader-like large cell and regular-size cell-mediated motion, whereas rotation emerges when large-cell-generated torque overcomes translation while cell-cell adhesion is maintained. Thus, collective migration mode depends on the balance among myosin-driven torque, regular-size cell-mediated translation, substrate coupling, and cell-cell adhesion. These modeling results suggest that collective epithelial migration can emerge *in vitro* from reciprocal biomechanical interactions between distinct cell states, rather than from leader-like cell behavior alone, and that such interactions can produce a predator-prey-like pursuit-escape mode of collective migration. Our model also provides a framework for investigating the biochemical and biomechanical regulation of collective cellular migration. It can be readily extended from *in vitro* to *in vivo* context and can incorporate the effects of substrate topology and soluble signaling factor-driven chemotaxis.

**Author summary:** When epithelial cells repair a wound, they move as coordinated groups rather than as isolated individuals. Cells of different sizes may contribute in distinct but connected ways. We developed a computational model to examine how a large cell and neighboring regular-size cells move together. In the model, cells attach to the underlying surface and one another, form protrusions that set their direction, and redistribute the force-generating protein myosin after contact. We found that group motion depends on a balance among surface attachment, cell-cell adhesion, movement of regular-size cells toward the large cell, and myosin-driven turning of the large cell. Depending on this balance, a mixed-size cluster can travel along a nearly straight path or rotate persistently, exhibiting pursuit-and-escape-like interactions between the large cell and surrounding regular-size cells. Rotation occurs when the large cell’s turning effect outweighs the translational motion driven by regular-size cells, while cell-cell adhesion keeps the cluster together. Our results show that collective migration can emerge from reciprocal mechanical interactions between cells in different states, not only from a “leader” cell acting alone. The model also provides experimentally testable predictions for how cell size, adhesion, and internal myosin distribution shape cell cluster movement.

## 1 Introduction

Wound closure, local cancer spread, and tissue-level processes during development rely on the ability of cells to migrate collectively, while maintaining contact with each other as well as coordinating their internal molecular- and organelle-level dynamics [1]. Compared with cells moving alone, collective cell migration has many advantages [2, 3]. For example, cell clusters exhibit enhanced sensitivity to biochemical and biomechanical signals, enabling them to respond to the surrounding cues more effectively [4]. Cell groups generally exhibit more persistent and directional migration than individual cells, which often display greater stochasticity and lower directional persistence. Collective migration can involve functional specialization among cells within a migrating group, allowing guidance, traction generation, and force transmission to be distributed across the collective rather than being borne by a single leader cell, thereby enhancing the robustness of collective movement [5, 6, 7, 8].

During wound healing, such as in the skin, collective migration of epithelial cells often involves two morphologically and, likely functionally distinct cell populations. Rare prominently large cells form at leading edges of migrating tissue sheets, while numerous regular-size cells trail behind. Large cells have flattened morphology and, thus, occupy a physically large area on the underlying substrate. In epithelial monolayer culture *in vitro*, visibly enlarged cells may reflect underlying differences with their regular-size counterparts in polarity, traction generation, cortical contractility, adhesion, cell-cycle state, or molecular signaling. Many of these features are poorly characterized experimentally, owning to challenges separating large from regular-size cells. We, therefore, used cell size as an observable marker of an assumed broader biomechanically and biophysically distinct cell state [9, 10, 11, 12]. Traditionally, large and regular-size epithelial cells have been referred to as “leaders” and “followers”, respectively, with the understanding that large leaders actively guide collective migration of remaining neighboring cells [9, 10, 13]. Indeed, *in vitro* studies using MDCK (Madin-Darby canine kidney) epithelial cells support this notion. Leader and follower MDCK cells emerge spontaneously in culture and form finger-like protrusions along the leading edge of a migrating cell sheet, while selective ablation of large cells impairs collective migration [14, 15]. However, recent studies showed that regular-size follower cells can also play active roles, challenging conventional “leader-follower” paradigm [16, 17]. For instance, during zebrafish gastrulation, it was shown that regular-size cells steer collective movement of the entire cell cluster by modulating speed and direction of cells directly ahead of them [18]. In zebrafish’s posterior lateral line primordium, regular-size cells can even push the primordium forward, complementing guidance exerted by large cells at the migration front [19].

Interactions between large and regular-size epithelial cells have been extensively studied in confluent cell monolayers *in vitro* [14, 5, 20]. In contrast, less is known about how large and regular-size cells interact within small discontinuous clusters [12]. Observations from different systems suggest multiple possible mechanisms, including large cell-driven pulling, regular-size cell-driven propulsion with large cells providing directional guidance, or cluster advancement resulting from regular-size cell proliferation [16]. Given the complexity of interactions within continuous epithelial sheets, examining smaller cell clusters provides a valuable opportunity to isolate and investigate fundamental mechanisms underlying collective cell migration.

In addition, collective migration of epithelial clusters composed of regular-size cells only has been studied *in vitro*, including investigations of molecular regulation of cytoskeletal dynamics at cell-cell junctions, lamellipodial activity, and galvanotactic responses [21, 22]. In particular, spontaneous rotational motion has been frequently observed in small epithelial cell clusters, including in cell doublets, in which two adherent cells rotate relative to each other while maintaining contact [23]. Several mechanisms and computational models have been proposed to explain this collective bahavior [24, 25, 26, 27]. Leong et al. used dissipative particle dynamics simulations to show that collective rotation can arise from front-to-rear asymmetry in paired cells, producing an S-shaped interface [24]. Camley et al. used phase-field model of cell motility to test various cell polarization principles and found that persistent rotational motion is most robust when cell polarity aligns with instantaneous velocity [25]. More recently, Lu et al. reported that spontaneous rotation of epithelial cell doublets is driven by a polarized myosin distribution, with myosin being recruited and activated near cell-cell contact interfaces and asymmetrically localized near the contact region [23]. Motivated by these findings, we implemented myosin-driven polarity determination mechanism into the modeling framework and investigated how different patterns of myosin redistribution can regulate motion of adhered regular-size cell pairs. We further tested this mechanism in mixed-size cell clusters composed of one large cell and several regular-size cells to assess its impact on collective motion.

Previously developed models for studying cell motility involved continuum methods, such as free boundary model [28] and phase-field model [29, 30], as well as lattice-based approaches, such as the Cellular Potts model [3, 31, 32]. Typically, computational models of cell motility on a substrate rely on the description developed by Abercrombie, in which cell movement results from dynamics of the actin network, myosin motors, and cell-substrate adhesion [33]. Subcellular element (SCE) modeling approach, where a cell is represented by a collection of interacting nodes, has been used to study the onset of individual cell movement [34]. The authors applied SCE model to study single cell crawling with a focus on how protrusions, contractions, and adhesion depend on intracellular chemical signals. Advanced SCE models have been also developed and applied previously to study blood clotting, early tissue morphogenesis and robust maintenance of tissue structure in plants [35, 36, 37]. The key advantage of the SCE framework is that its sub-modules have clear biological interpretations, enabling parameter calibration using experimental data and facilitating integration of biomechanical and biochemical effects.

In this work, we present a novel SCE model that incorporates both cell-substrate and cell-cell interactions to investigate collective motion of clusters composed either of two same-size adhering cells or of one large cell together with several regular-size cells. First, we investigated single-cell migration driven by cell-substrate interactions and protrusion-dependent polarity. Then, the impact of actomyosin dynamics in regular-size cells, represented by intracellular myosin levels, was studied to assess how different actomyosin dynamics influence the persistence or suppression of rotational motion in two-cell assemblies. Finally, we extended the analysis to small clusters containing one large cell and several regular-size cells, to test how myosin accumulation/activation, cell-substrate and cell-cell interactions can regulate collective migration of the entire cluster.

## 2 Results

### 2.1 Experimental observations identify contact-dependent large-cell turning in mixed-size clusters

We start with presenting the results of our analysis of primary experimental data in form of time-lapse videos capturing initial stages of interaction between regular-size MDCK cells and a large MDCK cell grown *in vitro* under pre-confluent, low-density starting conditions. Four representative examples are shown in Supplementary Videos A-D and in Figure 1. Below, we label these four video datasets as A, B, C, and D, corresponding to the four rows of Figure 1. In these representative video examples, contact between regular-size and large cells is accompanied by significant deformation of the latter, with its shape broadening in the direction away from the cell cluster. Examination of temporal motion dynamics by a large cell in response to regular-size cells suggests that flexible polarization strategy can allow for it to dynamically adjust its orientation to optimize the direction of movement.

**Figure 1:**
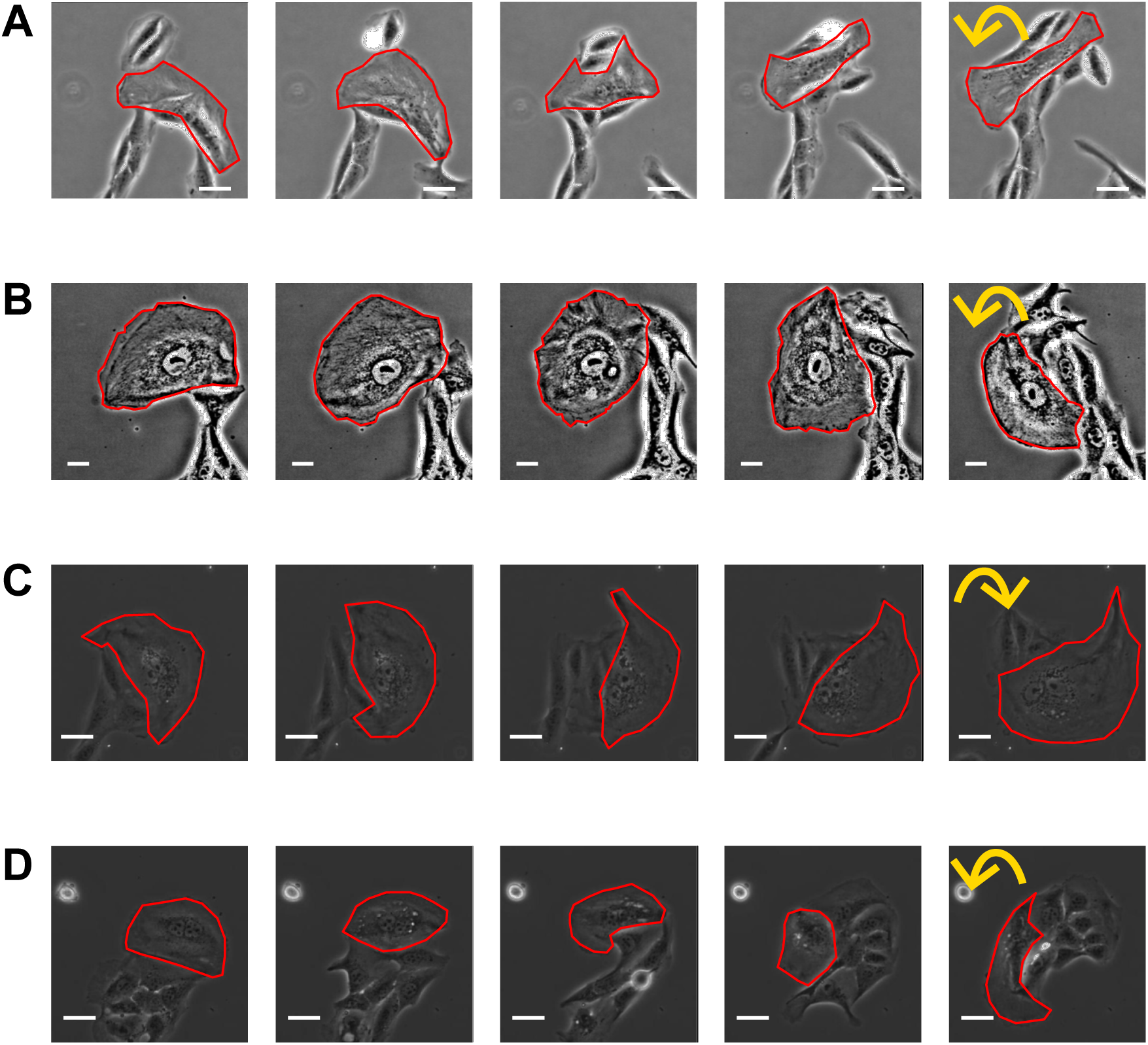
Representative experimental observations of interactions between a large cell and several regular-size cells. Each row corresponds to a single observation, recorded as time-lapse video. Time between consecutive snapshots is two hours. Contours of large cells (red) were manually extracted based on visual inspection of experimental images using ImageJ. Yellow semi-circular arrows depict directions of clockwise and counter-clockwise turns by large cells. Scale bar corresponds to 20 *µ*m.

#### Both large and regular-size cells begin to deform and move after establishing contact, resulting in coordinated motion

Upon their initial interaction, observable dynamics between large and regular-size cells resemble these of predator-prey behavior, where large cell acts as the prey and regular-size cells as predators. For example, in the dataset B (second row in Figure 1) all regular-size cells in the cluster appear to become active after one of them makes initial contact with a large cell, which is followed by them collectively chasing and surrounding it as if a prey. In datasets A, C, and D (as seen in the corresponding rows of Figure 1), already “chased” large cell “collides” with a second cluster of regular-size cells and, consequently, changes its direction to move away from both cell clusters. Datasets B and D, which capture large cell both before and after establishing contact with regular-size cells, indicate that large cell becomes triggered to move after making contact. To better depict this, in Figure 2 we traced the position of large cell nucleus in dataset B and the midpoint between two nuclei in dataset D (due to large cell being multi-nucleated). The resulting nucleus trace clearly shows that large cell exhibits random motion with minimal displacement prior to contact (black dots in Figure 2), while after contact its motion becomes faster and more persistent (green, blue, and red dots in Figure 2).

**Figure 2:**
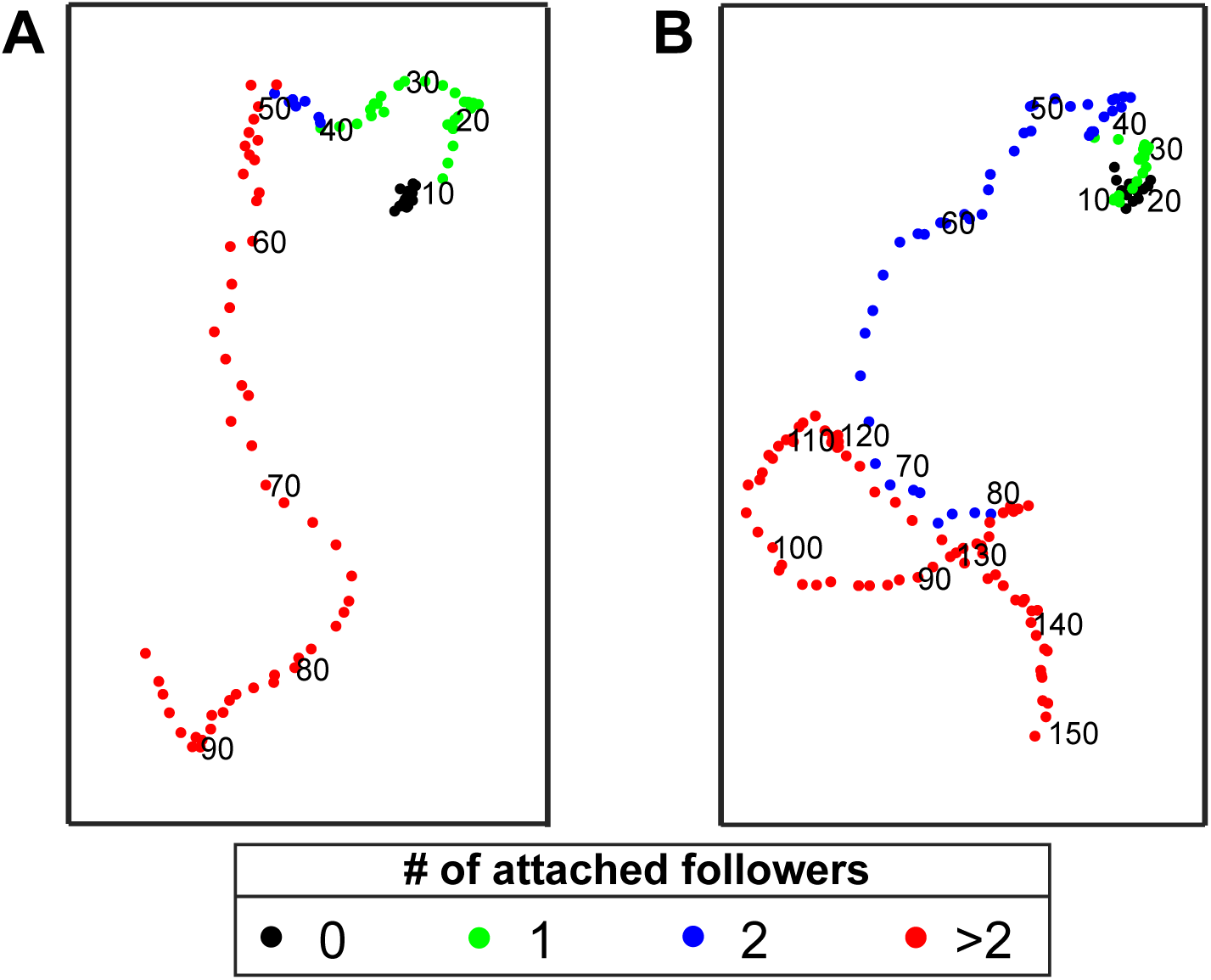
Displacement dynamics of large cells. Dots represent locations of large cell nucleus from datasets B (left) and D (right). Each dot corresponds to a single frame from a time-lapse video. Time between two sequential frames is 15 minutes. For every tenth frame, we indicate frame number near the corresponding dot. Color indicates the number of attached regular-size cells at a given time (frame) as define in the inset.

#### Large cells exhibit U-turn motion after contact with regular-size cells

Our observations on MDCK cells in low-density culture show that upon contacting a small cluster of regular-size cells (fewer than five cells), large cells deform and move in the direction parallel to contact line, defined as the portion of its observable boundary adjacent to the attached regular-size cells. Over time, such large cell exhibits a U-turn (see the rightmost column in Figure 1). This contrasts with how large cells behave when they are in contact with large clusters of regular-size cells or at the leading edge of scratched epithelial monolayers [12]. In these *in vitro* situations, large cells are said to guide regular-size cells toward empty spaces, which is believed to play an important role in wound closure. Additionally, in our observations we noted that after making a U-turn, which involves assuming diverse shapes, large cell eventually acquires stable crescent shape, with regular-size cells trailing behind. This is demonstrated in Figure 3, where bottom row shows contours of large cells before (black) and after (red) U-turn (datasets B, C, and D). In contrast, for the dataset A, regular-size cells contact large cell from multiple sides during U-turn, making its shape changes more complex (three representative contours are shown). We posit that once activated, a large cell acquires a stable shape, becomes polarized, and begins moving persistently, “pulling” a cluster of regular-size cells with it. To the best of our knowledge, this observation of a large cell briefly rotating with regular-size cells while establishing a distinct polarized direction, followed by persistent directional motion, has not been previously recognized as a feature of collective cellular dynamics of MDCK epithelial cells *in vitro*. Comparing these dynamics with previously reported persistent rotational motion of same-size MDCK cells [25, 38, 39, 40, 41, 42] suggests that addition of just one large cell is sufficient to result in a significant linear cluster displacements.

**Figure 3:**
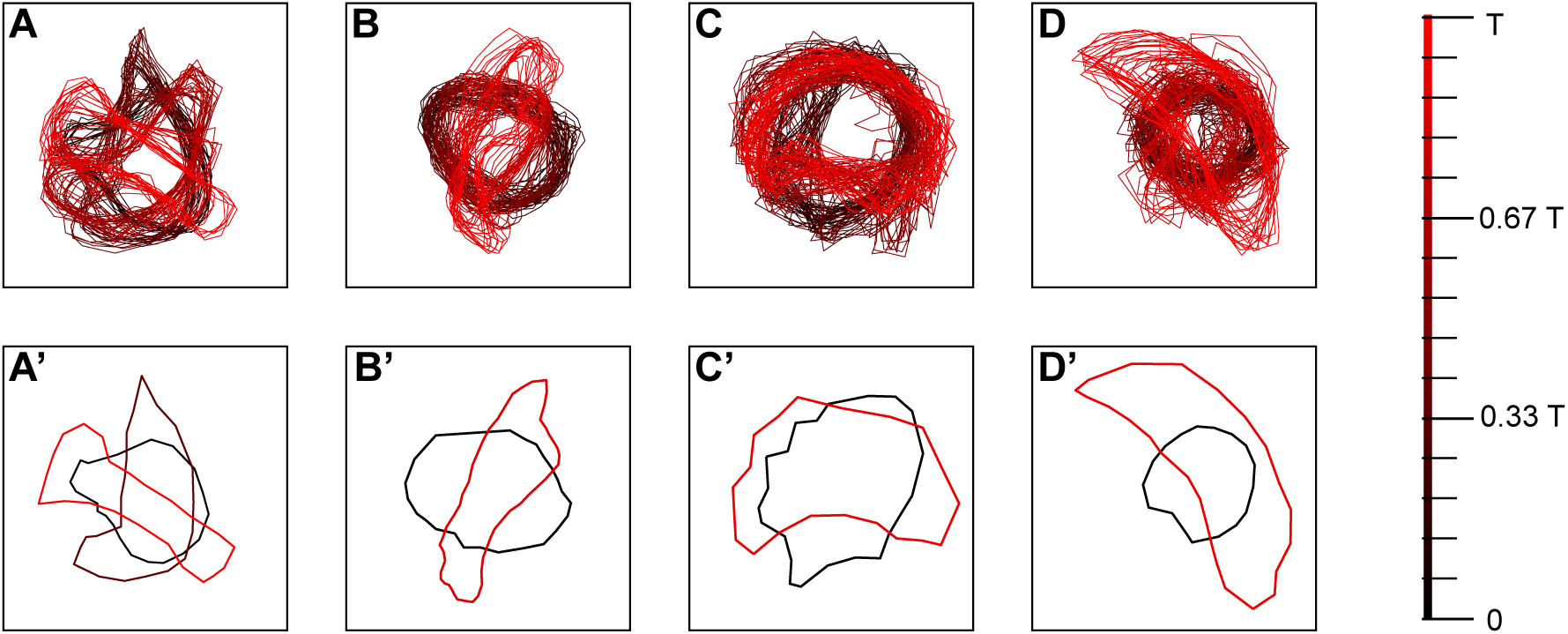
Dynamical shape changes by large cells. For datasets presented in Figure 1, contours of large cells are depicted in varying shades of red, with each shade corresponding to different time points. Correspondence between time points (here, *T* is the total observation time) and colors is depicted on the right. Bottom row shows representative contours of large cells.

#### Large cell moves in the direction parallel to the line of contact with regular-size cells

Next, we set out to validate that upon its activation, a large cell moves in the tangent rather than perpendicular direction with respect to contact line it forms with regular-size cell. To that end we computed large cell’s displacement directions and compared them to directions computed from the rule based on the assumption that it tries to move away from such contact line. The latter corresponds to either the “leading” role of a large cell in guiding follower cells to occupy empty space or to the predator-prey interaction, when large cell tries to “escape” from “chasing” follower cells. In either case one would expect large cell to move in the direction away from the contact line. Specifically, we designated *γ*_GL_ to be the curve describing such contact line.

Next, we computed the escape orientation by the path integral as follows:

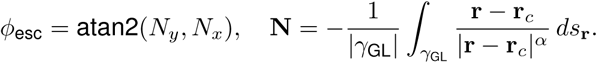

Here, large cell center **r***_c_* is defined as the center of its nucleus. Alternatively, in case of bi-nucleated large cell – as the midpoint of the segment connecting centers of two nuclei (see Figure 4A). Vector **N** = (*N_x_, N_y_*) points away from the contact line. Since the distance between point **r** on the contact line and cell center **r***_c_* varies along *γ*_GL_, vector **N** depends on the choice of the parameter *α*. However, value of *ϕ*_esc_ remains approximately constant across a wide range of *α*. Here, we chose to use *α* = 1. Finally, we computed *ϕ* as the angle between vector **N** and direction of displacement of the cell center. We present histograms for sin^2^ *ϕ* in Figure 4B-E for each dataset. Note that sin^2^ *ϕ* ≈ 0 corresponds to the motion of large cell toward or away from the contact line, whereas sin^2^ *ϕ* ≈ 1 corresponds to the motion parallel to the contact line. Histograms show that large cell mostly moves along the contact line in datasets A and D. Motion along the contact line over a substantial fraction of time is also visible in datasets B and C. These histograms suggest that large cell tends to move along the contact line in response to establishing contact with regular-size cells.

**Figure 4:**
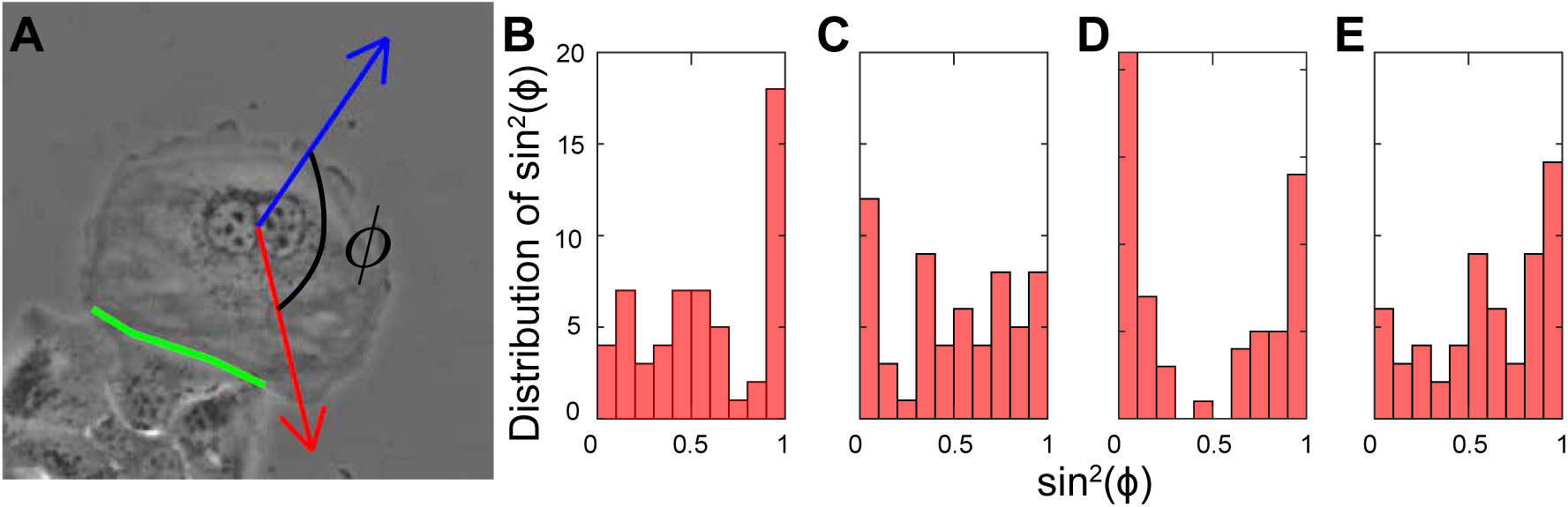
Large cell orientation dynamics. (A) Illustration of the angle *ϕ* between the direction of cell center displacement (red arrow) and the direction of motion away (blue arrow) from contact line (green line). (B-E) are histograms for sin^2^(*ϕ*) for datasets A through D presented in Figure 1.

Together, these observations provide three qualitative benchmarks for evaluating the mechanistic model. First, contact with regular-size cells should be associated with activation of large-cell displacement (Figure 2). Second, the model should be able to generate contact-dependent deformation of the large cell, including crescent-like shapes and U-turn-like reorientation (Figures 1 and 3). Third, be- cause experimentally observed displacement is often tangential to the contact line rather than purely normal to it (Figure 4), the model should allow contact geometry to redirect large-cell polarity away from a simple escape direction. We therefore used these observations to guide the development of a computational model in which contact-induced polarity changes, cell-substrate traction, and regular-cell pursuit determine whether a mixed-size cell cluster undergoes straight translation or sustained rotation.

### 2.2 A computational model links contact-induced polarity regulation to traction-driven cluster motion

Guided by the experimental observations shown above, we developed a multi-scale cell-based computational model (see Methods for details) for motile, interacting cells to investigate different modes of collective motion in regular-size cell doublets as well as in clusters consisting of one large cell and several regular-size cells. The model was calibrated using experimental data as described in the Supplemental Materials. Main model parameters are shown in Table 1. In this formulation, regular-size cells can maintain contact with the large cell through protrusion-based sensing and biased migration, while contact-induced myosin redistribution in the large cell provides a mechanism by which the contact interface can redirect large cell polarity. Cell-substrate adhesion parameters then control whether this polarity bias is converted primarily into translation or into torque of the entire cluster.

**Table 1:** Model parameters, their values and sources. Parameters labeled as “This study” represent effective parameters chosen to capture qualitative cellular behaviors in observed experiments.

| Parameter | Description | Value | Source |
| --- | --- | --- | --- |
| $\eta$ | damping coefficient for nodes dynamics | $3.6 \text{ nN} \cdot \text{min}/\mu\text{m}$ | This study |
| $\eta_a$ | damping timescale for polarity reorientation dynamics | 1 min | This study |
| $N_{\text{adh}}^{\text{max}}$ | maximum number of cell-substrate adhesion sites allowed per node | 1 – 10 | This study |
| $R_r$ | initial radius of a regular-size cell | $15 \mu\text{m}$ | [12, 83] |
| $R_l$ | initial radius of a large cell | $50 \mu\text{m}$ | [12, 84] |
| $N_{\text{prot}}$ | maximum number of simultaneous protrusions per cell | 5 | This study |
| $L_{\text{max,p}}$ | maximum extension length of a protrusion | $5 \mu\text{m}$ | [74, 85] |
| $T_p$ | lifetime of a protrusion | 2 – 10 min | [86, 87, 88] |
| $P_p$ | rate of protrusion initiation for a regular cell | $0.0137 \text{ min}^{-1}$ | This study |
| $k_p$ | effective reorientation rate induced by protrusion activities | $0.054 \mu\text{m}^{-1}$ | This study |
| $k_a$ | amplification factor representing enhanced polarity coupling upon protrusion-cell sensing | 2 | This study |
| $k_l$ | effective polarity alignment rate toward the direction of a large cell | 3 | This study |
| $\mu_{\text{adh}}$ | mean initial length of a focal adhesion | $1.845 \mu\text{m}$ | [81] |
| $\sigma_{\text{adh}}$ | standard derivative of the focal adhesion length distribution | $0.513 \mu\text{m}$ | [81] |
| $k_{\text{SA}}$ | effective spring constant governing cell-substrate adhesion | $0.006 - 0.42 \text{ nN}/\mu\text{m}$ | This study |
| $k_{\text{on}}$ | basal binding rate of focal adhesion formation | $0.4 - 2 \text{ min}^{-1}$ | This study |
| $k_s$ | characteristic area scale controlling adhesion binding probability | $5.56 \times 10^2 \mu\text{m}^2$ | This study |
| $l_0$ | characteristic rupture length of a cell-substrate adhesion bond | $1.67 - 3.33 \mu\text{m}$ | This study |
| $m_0$ | characteristic myosin level at which cell-substrate adhesion rupture occurs | 0.5 | This study |
| $d_{\text{tr},1}$ | threshold distance from the contact interface required to initiate myosin recruitment | $13.3 \mu\text{m}$ | This study |
| $d_{\text{tr},2}$ | characteristic averaging distance used to compute membrane-associated myosin levels | $13.3 \mu\text{m}$ | This study |

#### Impact of cell-substrate adhesion and protrusion-dependent polarity on individual cell motility

First, we simulated dynamics of individual cells to demonstrate that the model adequately captures autonomous cell motion. One of the key components governing cell motility is cell-substrate adhesion. We studied how dynamic cell-substrate adhesion regulates cell motility by balancing traction generation and adhesion turnover. Specifically, we analyzed how individual cell’s speed depends on parameters governing cell-substrate adhesion, including (i) substrate adhesion coefficient *k*_SA_, (ii) characteristic unbinding length of adhesion bonds *l*_0_, (iii) baseline adhesion formation rate *k_on_* and (iv) maximum number of adhesion sites that can be formed at a single node *N* ^adh^ . The results are shown in Figure S1. We further quantified single cell motility by calculating mean squared displacement (MSD) as a function of time lag *τ* . The model was able to capture key qualitative features of isolated cell movement. Mean-squared displacement analysis revealed ballistic behavior at short time lags, in which MSD grows approximately as *τ* ^2^, and displays super-diffusive or diffusive behavior at longer times (Figure S2) [43, 44]. We further characterized how the diffusion coefficient depends on adhesion parameter *k*_SA_, cell protrusion extension rate *P_p_* and protrusions lifetime *T_p_* (Figure S3). This approach provided a mechanistic framework that directly links underlying cellular processes to anomalous (non-Brownian) translational diffusion observed across diverse cell types, including CD8^+^ T cells during target search and invasive cancer cells during tumor metastasis [45, 46, 47], thereby enabling systematic investigation of the mechanisms driving these broadly present cellular behaviors.

#### 2.2.1 Asymmetrical myosin accumulation induces rotational motion in a pair of adhered cells

Next, we extended the model to simulate cell doublets consisting of two adhering regular-size cells. The key additional feature of the model was that on top of mechanical interactions between cells via cell-cell adhesion and volume exclusion, cell-cell adhesion was assumed to regulate myosin distribution within both cells and consequently, their polarity. This is consistent with experimental observations showing that cadherin-dependent adhesions modulate spatially polarized actomyosin contractility and influence front-to-rear polarity in collective cell migration [48, 49, 50, 51, 52]. Motivated by experimental observations of polarized myosin accumulation [23], we defined two contact response parameters *γ*_sym_ and *γ*_asym_ (see also Eq. (3) in Methods), which quantify rates of uniform and polarized myosin recruitment toward contact interface, respectively. In the following, we focused on three representative cases: *symmetric flux* (*γ*_asym_ = 0), *mixed flux* (*γ*_asym_*/γ*_sym_ = 1), and *asymmetric flux* (*γ*_sym_ = 0). The resulting myosin distributions determined each cell’s polarity direction and, as the result, governed their subsequent motions following the initial contact. Depending on the myosin profile, a cell doublet may exhibit either short-lived or persistent rotation.

As the initial condition for our simulations, two cells were initially positioned close to each other, so that cell-cell adhesion bonds were formed almost immediately. Figures 5A-C show the initial configurations of cell doublets, along with trajectories of their geometric centers, defined as the average location of all nodes representing each cell. To assess whether a doublet exhibits persistent rotation, we tracked the midpoint between two cell centers and the displacement of each center relative to this point. Persistent rotation corresponds to a sustained angular velocity of cell centers. Simulation results showed that persistent rotation arises only under asymmetric myosin flux, in which myosin was recruited from other regions of the cell toward nodes on one side of the contact interface, producing polarized myosin localization. Specifically, as shown in Figure 5B, both cell centers followed circular trajectories around their midpoint. In contrast, under symmetric myosin flux condition (Figure 5A), myosin was recruited from other regions of the cell toward all nodes within a predefined distance *d*_tr,1_ of the contact interface, resulting in an approximately uniform distribution along the interface. In this case, rotational motion of cell doublet was suppressed. When mixed flux condition was considered, in which symmetric and asymmetric components superimposed (Eq.(3)), rotational dynamics were sustained but weakened, with reduced angular velocities compared with the asymmetric case (Figure 5C).

**Figure 5:**
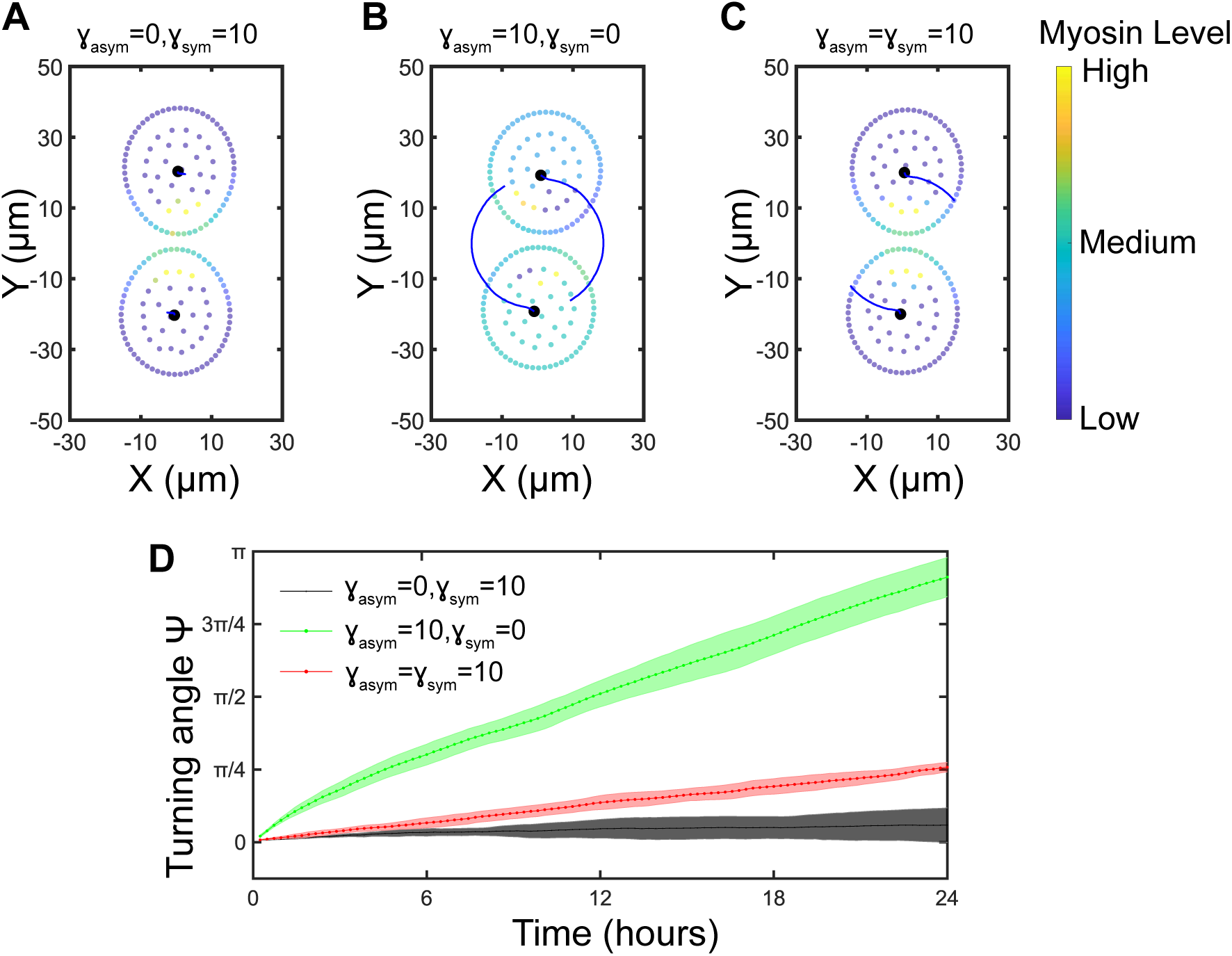
Simulation results for two adhered regular cells. (A,B,C) The initial configuration of two adhered regular-size cells, each represented by its membrane and internal nodes for three myosin flux cases: symmetric (A), asymmetric (B), and mixed (C). To characterize trajectories of the cells, we compute their geometric centers defined by the average *x*- and *y*- coordinates of all nodes (both internal and membrane) of the corresponding cell. Initial locations of geometric centers of the two cells are shown. (D) Turning angle *ψ* is defined as the angle between the segments connecting the geometric centers of two adhered cells at the initial and current time points. Results are shown for three cases: *γ*_asym_*/γ*_sym_ = 0 (asymmetric), = 1 (mixed), and = *∞* (symmetric).

These results can be interpreted as follows. The model assumes that each cell’s polarity vector points away from the region of highest myosin accumulation, so that when myosin is concentrated near cell-cell contact, polarity is biased toward the side with less myosin. In the asymmetric flux case, myosin concentrates at one end of the contact in one cell and at the opposite end in the other cell, so that each cell’s polarity vector is rotated away from the line joining two cell centers. Adhesion bonds to the substrate therefore transmit traction forces along those rotated polarity directions and, combined with cell-cell adhesion, they generate a net torque that drives persistent doublet rotation. By contrast, under symmetric flux, myosin distribution is nearly uniform along the contact interface. The two cells therefore polarize approximately away from the shared interface in opposite directions, so that the resulting substrate traction forces largely oppose one another rather than forming a persistent torque. As a result, the doublet remains mechanically stable rather than rotating. For the mixed flux case, the outcome depends on the relative strength of the asymmetric versus symmetric components. If the asymmetric component dominates, myosin maximum shifts to one side of the interface and rotation persists. If the symmetric component dominates, myosin maximum stays near the interface midpoint and rotation is attenuated. Figure 6A shows the myosin profiles centered on the contact midpoint for symmetric, asymmetric and mixed cases. Figure 6B shows quantified asymmetric myosin distribution along the contact using asymmetry index 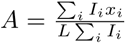 where *I_i_* is myosin intensity at membrane node *i*, *x_i_* is the membrane node position indexed relative to the contact center, and *L* = max(|*x_i_*|) is the normalization constant. The asymmetry index measures polarization of myosin distribution along the cell-substrate contact interface and quantifies whether myosin is biased toward one side of the contact center. Values range approximately from −1 (all myosin is at one end) to +1 (all myosin is at the other end), with 0 indicating symmetric distribution. Figure 6C shows corresponding net torque generated by cell-substrate forces at membrane nodes, computed as *τ*_net_ = *Σi* [(*x_i_* − *x_c_*)*F_y,i_* − (*y_i_* − *y_c_*)*F_x,i_*], where (*x_c_, y_c_*) denotes doublet center. Symmetric case fluctuates around small torque, whereas asymmetric case develops sustained torque in the direction prescribed by the imposed myosin asymmetry. Mixed case shows intermediate response, consistent with a weaker force imbalance than in fully asymmetric case. We also plotted dependence of angular velocity of doublet rotation in the asymmetric case on adhesion stiffness (*k_SA_*). In Figure 6D angular velocity initially increases with (*k_SA_*) in a saturating, Hill-like fashion, which is consistent with the increase in single-cell motility (also see Supplement Figure S1). As adhesion strength increases further, angular velocity decreases, and doublet splits in the case of high adhesion stiffness. This may arise because stronger cell-substrate adhesion amplifies traction component along the cell-cell axis, pulling two cells apart and thereby disrupting coordinated rotation of the doublet.

**Figure 6:**
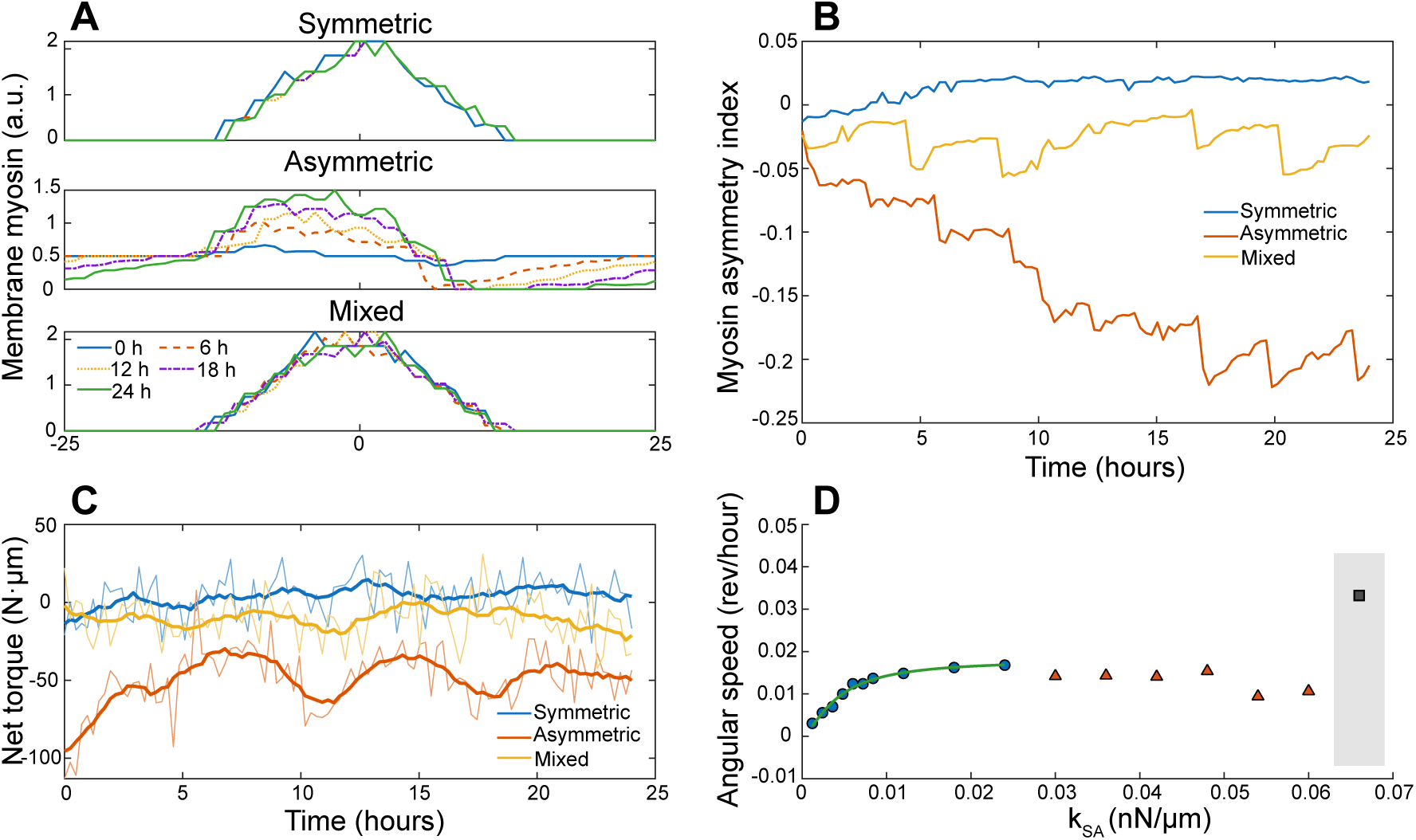
Biomechanical and biochemical coordination in cell doublets under three different mechanisms of myosin recruitment and adhesion strength. (A) Membrane myosin intensity profiles for symmetric, asymmetric, and mixed cell doublets at selected time points (0, 6, 12, 18, 24 h) across membrane nodes numbered from -25 to 25, such that node 0 corresponds to the midpoint of the contact line. (B) Dynamics of the myosin asymmetry index *A*, computed as the first spatial moment of the membrane myosin profile normalized by total myosin intensity and cell membrane length. (C) Net torque around the doublet center generated by membrane force (light lines) and their smoothened counterparts (thick lines) for the three cases of membrane myosin intensity profiles. (D) Angular velocity while adhered as a function of the adhesion stiffness coefficient (*k_SA_*), with low-(*k_SA_*) points fit by a Hill-like response. Shaded region indicates doublet separations.

Notably, our simulations predicted angular velocity in the range of approximately 0.01−0.02 rev/hour, which is less than experimentally measured velocity of 3D epithelial cell doublets rotation (0.2 − 0.5 rev/hour) or that of HUVECs (Human Umbilical Vein Endothelial Cells) on disk-shaped micro-patterns, measuring 1 − 2 rad/hour (0.159 − 0.318 rev/hour) [23, 38]. Within the parameter ranges considered here, changes in parameters, such as substrate adhesion coefficient, did not significantly alter predicted magnitude of angular velocity as long as the system remained in the rotational mode. However, it is known that patterned substrates tend to promote consistent rotation [38]. On patterned dishes, pairs of human epidermal keratinocytes can rotate at angular velocities less than 0.0625 rev/hour, which is in a range comparable to that predicted by our simulations [53]. Overall, our simulations show that asymmetrically localized myosin is capable of driving rotational motion in cell doublets, when cells are similar in size and cell polarity is defined to point away from the region of highest myosin concentration [23]. Furthermore, as large cells can spontaneously arise within epithelial colonies *in vitro*, it is important to investigate how such large cells respond to contact-induced myosin redistribution and how this affects movements of heterogenous cell clusters.

#### 2.2.2 Mixed-size clusters reproduce contact-dependent translation and rotation

Next, we simulated dynamics of a large cell interacting with a group of regular-size cells. As in the cell doublet simulation, large cell and regular-size cells were initially placed close together, so that contact can be established quickly (Figure 7). Specific initial arrangement of regular-size cells resembled that in our experimental observation from the dataset B, shown in Figure 1. To isolate the response by a large cell to contact with regular-size cells, we assumed that polarity of each regular-size cell was determined by the combined effect of stochastic protrusion extensions, following the same mechanism as described for isolated single cells. This assumption represents a simplification of the intracellular dynamics of regular-size cells. Specifically, regular-size cells were treated as actively motile units whose motion is biased toward large cell, thereby ensuring stable attachment and cluster formation. Within this framework, we did not attempt to resolve detailed intracellular dynamics of regular-size cells, nor did we include feedback effects on their motility.

**Figure 7:**
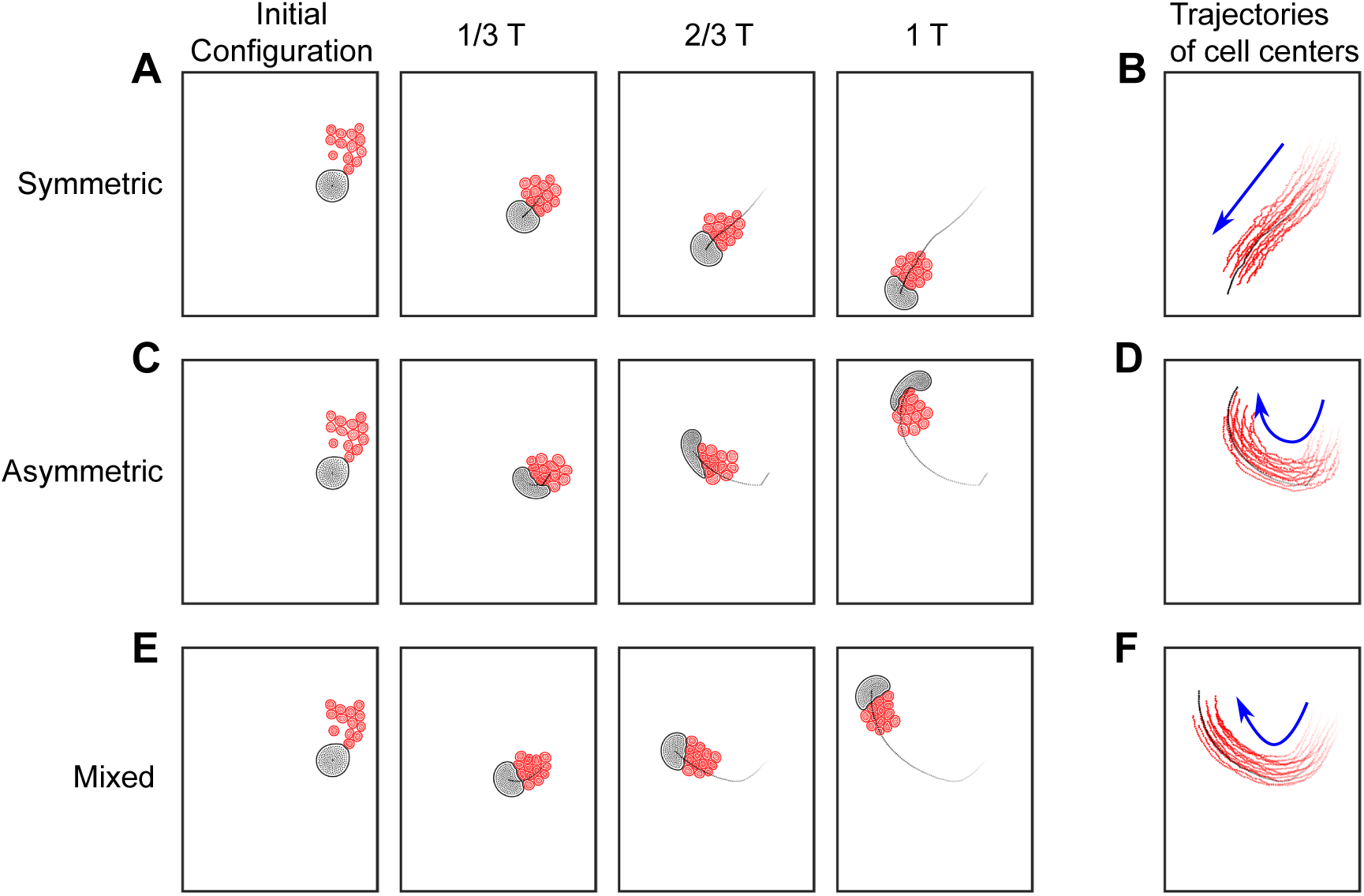
Simulations of clusters composed of a large cell and several regular-size cells for different myosin flux cases. Each of the three rows represents specific myosin flux case: symmetric (A,B), asymmetric (C,D), and mixed (E,F). Regular-size cells are depicted in red, and large cell is shown in black. For each case, the first four panels (A,C,E) show cluster configuration at times near 0, ^1^ *T*, ^2^ *T*, and *T*, where *T* denotes full simulation time. Each panel also shows the trajectory of the geometric center of large cell up to the time point depicted. Right-most panels (B,D,F) show the complete trajectories of individual cells: regular-size cell are shown with red tracks, and large cell with black track. Trajectory intensity decreases with time to indicate temporal progression. In addition, a blue arrow illustrates the overall direction of motion.

In addition, regular-size cells were biased to migrate toward a neighboring cell when the neighboring cell resided within the sensing range of an extended cellular protrusion. Specifically, this occurred when the distance between the protrusion tip and the center of the neighboring cell was smaller than the neighbor’s radius, thereby enabling cell-cell aggregation. Furthermore, experimental observations suggest that visibly enlarged or leader-like cells in epithelial colonies can be mechanically distinct, exhibiting enhanced traction generation and stronger cell-substrate adhesion signaling [11, 14, 12]. Regular-size cells are known to respond to mechanical cues within tissues and can bias their migration toward regions of elevated tension or stiffness, which are frequently associated with mechanically dominant cells [54, 55]. In our model, this effect was incorporated phenomenologically by introducing a directional bias in the motion of regular-size cells toward a large cell. This modeling choice facilitated formation of initial cluster and promoted sustained interactions between regular-size cells and a large cell.

Like with same-size cell doublet simulations, we investigated three types of myosin redistribution in a large cell: symmetric, asymmetric, and mixed fluxes. We first examined whether large cell exhibits distinct migratory patterns upon attachment of regular-size cells with these three redistribution patterns. As depicted in Figure 7, regular-size cells first gathered together due to protrusion-based sensing and moved toward large cell due to imposed directional bias. Upon contact with regular-size cells, large cell first underwent significant shape changes, adopting crescent-like morphology with concave side facing regular-size cells. Due to directional bias introduced in regular-size cells, their migration was consistently oriented toward a large cell. Under symmetric myosin flux, polarity of a large cell aligned closely with the direction of regular-size cell motion and pointed away from the large-to-regular cell contact interface. As a result, the entire cluster exhibited persistent directional migration, driven either by leader-like motion of the large cell or by regular-size cell-mediated translation arising from their pursuit of the large cell, depending on the relative magnitudes of traction forces. In the case of asymmetric myosin flux, myosin accumulated primarily on one side of the large-to-regular cell contact interface, causing polarity of the large cell to shift toward the opposite side of the interface. This polarity shift generated a tangential force component and drove sustained rotation of large cell together with attached regular-size cell cluster. Because cell-cell adhesion maintained cluster cohesion, regular-size cells continued to pursue large cell while being carried into the rotational motion. In the mixed-flux case, location of myosin peak at different times depended on the strength of asymmetric and symmetric components. Thus, diverse motion patterns of a large cell and, consequently, that of the entire cell cluster, evolved as shown in Figure 7.

These simulations capture several qualitative features observed experimentally: (i) regular-size cells remain associated with large cell, (ii) large cell deforms at the contact interface, and (iii) the resulting cell cluster can transition between turning and directional migration. In the model, turning motion arises when asymmetric contact-induced myosin redistribution redirects large-cell polarity away from the normal escape direction while regular-size cells continue to pursue large cell. This mechanism provides a possible explanation for the experimental observation that large-cell displacement is often tangential to the contact line (Figure 4).

#### 2.2.3 Substrate coupling and large-cell adhesion capacity tune the translation-rotation balance

We also studied how the behavior of cell cluster depends on cell-substrate adhesion, which directly impacts individual cell motility. Adhesion forces in the model are directly regulated by two parameters: substrate adhesion coefficient (*k*_SA_) and maximum number of adhesion bonds a given cell can form (*N_l_* for large cell and *N_r_* for regular-size cells). Each adhesion site generates traction force toward the polarity of a cell. Increasing substrate adhesion strengthens force generated by each bond, while increasing the number of bonds adds traction forces parallel to cell polarity. Thus, both parameters are able to enhance cell motility (Figure S1). Under the condition of asymmetric myosin flux, we varied both the substrate adhesion coefficient *k*_SA_ and maximum number of adhesion sites *N_l_* in a large cell, and tracked trajectory of its geometric center. Maximum number of adhesion sites in regular-size cells *N_r_* was held fixed.

Representative trajectories of a large cell for different values of (*N_l_*) are shown in Figure 8. Once regular-size cells attached to large cell, cell-cell adhesive forces maintained cluster cohesion, allowing cells to migrate collectively. Therefore, trajectory of large cell center provided a representative measure of the motion for the entire cluster. When large cell formed relatively few substrate adhesion sites, its motility was low and cluster motion was dominated by regular-size cells. Because regular-size cells polarized toward large cell, this generates regular-size cell-mediated translational component that drove the entire cluster along nearly straight trajectory, even when asymmetric myosin flux was present in large cell.

**Figure 8:**
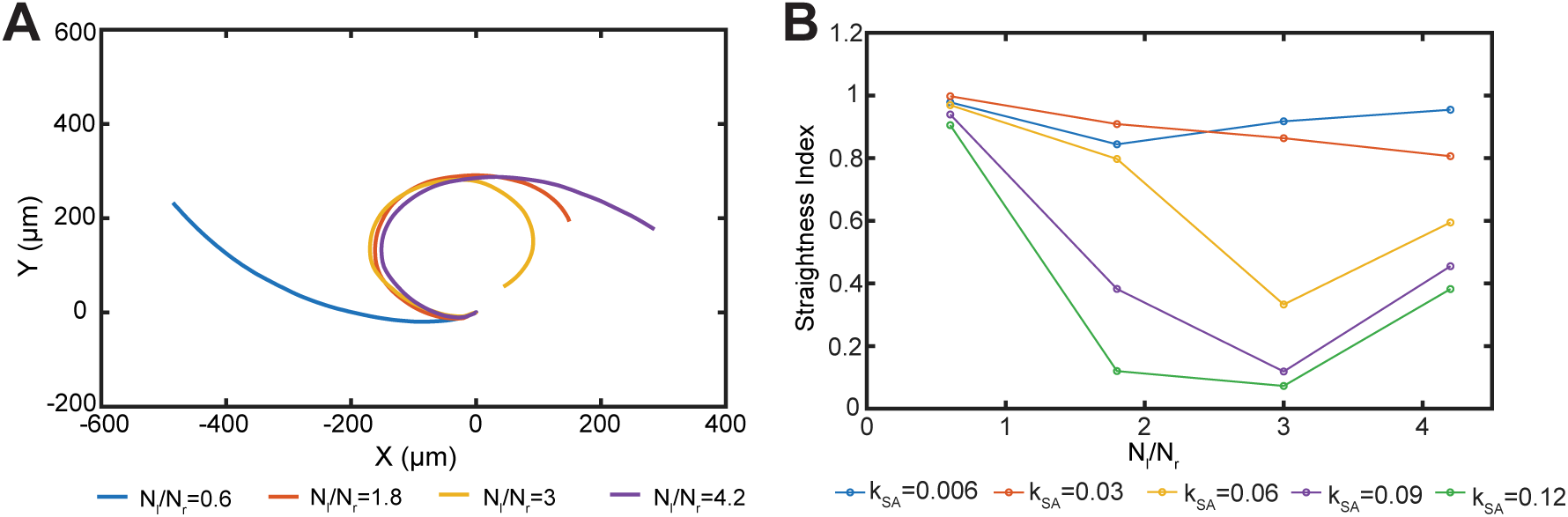
Dependence of migratory patterns of clusters containing one large cell on cell-substrate adhesion parameters. (A) Representative trajectories of a large cell for different ratios of maximum numbers of focal adhesion sites in a large cell and in a regular-size cell, with fixed substrate adhesion coefficient *k*_SA_(*nN/µm*) . (B) Dependence of the straightness index (SI) on the ratio of maximum number of total adhesion sites in a large cell relative to a regular-size cell, shown for different values of the substrate adhesion coefficient.

In contrast, within the parameter range examined here, increasing the number of substrate adhesion bonds formed by the large cell enhanced its effective traction generation and allowed it to become the primary driver of collective cluster movement [56, 57]. Under these conditions, asymmetric myosin flux caused large cell to repolarize away from contact-associated myosin-enriched region, generating a tangential force component. Together with the continued pursuit of large cell by regular-size cells and maintenance of cluster cohesion by cell-cell adhesion, this pursuit-escape interaction produced predator-prey-like rotational motion of the entire mixed-size cell cluster.

To characterize large cell trajectory, we computed straightness index (SI) for each trajectory, defined as:

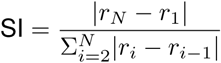

where *r_i_* denotes position of the geometric center of a large cell at time step *i*. The numerator represents net displacement of a large cell, and the denominator is the total path length. SI provides a measure of trajectory straightness and is often interpreted as an indicator of migration directionality. It has been widely applied in the analysis of solitary cell and cell cluster migration across various biological systems [45, 58, 59]. SI value being close to 1 indicates a nearly straight trajectory, whereas highly curved or rotational paths correspond to SI ≪ 1. Simulation results show that SI decreases as the relative adhesion capacity of large cell, *N_l_/N_r_*, increases (Figure 8B). For a fixed substrate adhesion coefficient, increasing maximum number of adhesion sites in the large cell enhanced traction generation and promoted rotational motion. Likewise, at fixed adhesion-site number, increasing substrate adhesion coefficient strengthened large cell traction, again favoring rotational motion of the cluster. Interestingly, dependence of SI on (*N_l_/N_r_*) was non-monotonic. For higher values of substrate adhesion coefficient, SI decreased as (*N_l_/N_r_*) increased up to approximately 3, but then increased again when (*N_l_/N_r_*) exceeded 4 (Figure 8B). This increase did not indicate a loss of rotational tendency, but rather reflected a change in cell morphology and the geometry of rotation. When (*N_l_/N_r_* ≤ 3), migration speed of large cell was comparable to that of the attached regular-size cells. Under these conditions, cell-cell adhesion and collective motion deformed large cell into a crescent-like shape. Because large cell polarity was defined relative to the contact-associated myosin distribution and distal membrane geometry, this deformation reduced the effective radius of rotation and produced a more curved trajectory of the large cell center. In contrast, when (*N_l_/N_r_ >* 4), large cell migrated more rapidly and underwent less pronounced deformation. The resulting polarity geometry generated larger effective rotation radius, causing large cell center trajectory to appear straighter over the simulated time window and thereby increasing SI.

Our analysis of clusters containing one large cell and several regular-size cells shows that spatial distribution of myosin within the large cell can determine the overall migratory behavior of the cluster when large cell is sufficiently motile to act as the primary motility driver. Relative motility of large cell and regular-size cells controls whether asymmetric myosin redistribution produces directed translation or predator-prey-like rotation. When large cell is minimally motile, motion of regular-size cells toward it dominates cluster displacement. In this regime, large cell contributes less to propulsion, so cluster displacement is dominated by the collective motion of regular-size cells toward the large cell, producing an almost straight trajectory despite asymmetric myosin flux in the large cell. In contrast, when large cell is highly motile, asymmetric myosin enrichment along large-regular cell contact interface introduces a tangential component to large cell polarity. Because regular-size cells continue to polarize toward large cell, system forms a predator-prey-like pursuit-escape interaction, with regular-size cells pursuing and large cell escaping. Under these conditions, large-cell-generated torque can dominate over regular-size cell-mediated translational component, leading to coherent rotation of the entire cluster while cell-cell adhesion maintains cohesion. Together, these results provide a mechanistic interpretation of how mixed-size clusters with the same asymmetric myosin redistribution in the large cell can switch between straighter displacement and contact-induced turning or rotation. In the model, increasing large-cell substrate coupling or adhesion capacity strengthens large-cell traction and allows the asymmetric polarity cue to drive rotational motion, whereas weaker large-cell traction allows regular-size cell-driven translation to dominate, leading to straighter collective displacement. Thus, our model identifies large-cell mechanical dominance as a control parameter that converts the same asymmetric myosin cue into distinct multicellular migration modes, providing testable predictions for how substrate adhesion and myosin organization regulate cluster dynamics.

## 3 Discussion

Understanding how epithelial cells coordinate their motion in small groups remains a fundamental challenge in the study of collective cell migration. While extensive work has examined collective migration in large continuous epithelial sheets, much less is known about how mechanical interactions and polarity dynamics govern behavior of small cell clusters during early stages of tissue organization, such as at the onset of wound healing. Here we developed a multiscale cell-based computational model that integrates cell-cell adhesion, cell-substrate adhesion, protrusion-based cell-cell interactions and polarity regulation to study mechanisms of migration in small epithelial cell assemblies. Model simulations tested several alternative mechanisms driving characteristic behaviors observed experimentally, including anomalous diffusion in single-cell migration, persistent rotational motion in cell doublets, and rotational and directed migration in heterogeneous mixed-size cell clusters.

Simulations of isolated cells highlighted important role of cell-substrate adhesion in regulating migration. We found that cell velocity in the model increases with substrate adhesion coefficient, baseline adhesion formation rate and maximum number of adhesion sites per node, whereas it decreases as characteristic unbinding length of adhesion bonds increases (Figure S1). Mean squared displacement analysis further showed that isolated cells exhibit ballistic motion at short time scales followed by normal or super-diffusive behavior at longer time scales rather than random walk, which is consistent with previously reported experimental observations [43, 44]. Such anomalous diffusion reflects active cellular motility rather than passive Brownian motion and is commonly associated with directional persistence during cell migration. Previous theoretical studies have shown that persistent random walks can reproduce similar statistical signatures [60, 61]. Our model complements these approaches by providing mechanistic framework in which anomalous migration behavior emerges from explicit representations of adhesion dynamics and polarity regulation through protrusion formation.

For cell doublets composed of regular-size cells, we implemented in the 2D SCE framework a previously proposed mechanism for persistent rotational motion that has been reported in several experimental systems [38, 62, 63]. Myosin activity is often enriched near cell-cell contacts, yet how this myosin localization influences collective motion remains unclear. Motivated by recent experiments showing that asymmetric myosin localization can drive persistent rotation of cell doublets [23], we implemented a polarity rule in which polarity vector points away from the region of maximal myosin accumulation. Simulations showed that asymmetric myosin distribution along cell-cell interface led to persistent rotation, whereas symmetric distribution suppressed it (Figure. 5). Moreover, angular velocity showed a non-monotonic dependence on substrate adhesion strength. At low adhesion strengths, it increased with increasing adhesion, consistent with enhanced traction facilitating the conversion of polarity-driven motion into rotational motion. At higher adhesion strength, however, angular velocity slightly decreased, and excessive adhesion led to cell separation rather than coherent rotation (Figure 6). Together, these results indicate that asymmetric myosin localization at cell-cell interfaces may act as a polarity cue that promotes persistent rotational dynamics in small cell groups.

We next examined clusters consisting of one large cell interacting with several regular-size cells. Large cells located at the leading edge of continuous epithelial sheets have traditionally been described as functional “leaders”, and their presence was thought to be essential for coordinated epithelial migration during wound healing [14]. However, recent studies suggest that interactions between large and regular-size cells are more complex than previously assumed, and that regular-size cells can actively contribute to driving collective migration [16, 27, 64]. Because of the spontaneous emergence of large cells and variability of their spatial arrangements, it is difficult to study early-stage cell cluster formation experimentally. Therefore, computational modeling provides useful approach for exploring potential interaction and clustering mechanisms.

Our model simulations revealed that heterogeneous-size cell clusters could exhibit qualitatively different migration patterns, including persistent rotation and directed motion, depending on the distribution of myosin within a large cell and relative levels of cell-substrate traction forces among all participating cells (Figure 7). When large cell generates stronger traction forces, it tends to guide migration direction of the cluster. Conversely, when regular-size cells collectively contribute a stronger translational component, they can drive motion of the entire group (Figure 8). These results suggest that, in our model, migration direction of heterogeneous-size cell clusters can emerge from a balance of biomechanical activity among different cells, rather than being solely determined by a fixed leader-follower hierarchy.

Our model suggests a non-canonical predator-prey-like mechanism for mixed-size epithelial cell cluster motion, and non-“leader” behavior for large cells. In our simulations regular-size cells actively polarized toward large cell. Polarity of the latter was determined by spatial distribution of contact-associated myosin enrichment: in both symmetric and asymmetric cases, large cell polarized away from myosin-rich region, but the geometry of this region determined whether the resulting motion was translational or rotational. When myosin enrichment was symmetrically distributed along large-regular-size cell contact interface, large cell polarity remained aligned with the overall direction of cluster displacement, producing near-straight collective migration. When myosin enrichment was biased toward one side of the contact interface, the resulting polarity acquired a tangential component and generated large cell torque. Together with the continued polarization of regular-size cells toward large cell, this produced a pursuit-escape-like interaction that could drive coherent rotation when large-cell-generated torque dominated over regular-size cell-mediated translation and when cell-cell adhesion maintained cluster cohesion. These results suggest that reciprocal polarity regulation between heterogeneous cells provides an additional biomechanical mechanism for collective migration in heterogeneous epithelial cell clusters.

While current model provides several important biologically-relevant insights, several simplifying assumptions were made. Simulations were performed in 2D and, therefore, did not capture the full 3D organization of epithelial tissues. In addition, cell polarity was represented in the model by simplified protrusion-dependent or myosin-dependent mechanisms, and cell-substrate adhesion dynamics were described using coarse-grained parameters. While these simplifications allowed systematic exploration of collective cell behaviors, incorporating more detailed biochemical signaling pathways and molecular interactions will be important in the future for testing different biological mechanisms and comparing test results with multi-scale experimental data. At the same time, the developed framework provides a flexible platform for extending the study of collective cell migration. In migrating epithelial cells, front-rear polarity is controlled by multiple signaling pathways, including Rho-family GTPases Rac1 and RhoA. Integrating these signaling networks into the model would enable investigation of how biochemical polarity circuits couple with biomechanical processes during collective cell migration. In particular, incorporating RhoA-ROCK signaling, which regulates actomyosin contractility, focal adhesion maturation, and traction force generation, may help clarify how intracellular signaling translates polarity cues into biomechanical force production.

Another promising future direction is the study of biochemically guided migration. In the present model, polarities in isolated cells and in regular-size cells within mixed clusters were determined by randomly oriented protrusions. In the chemotactic contexts, however, protruding structures, such as filopodia, play assisting role in sensing extracellular signaling molecule gradients. Chemokine gradients can regulate both the orientation and spatial distribution of filopodia, enabling cells to interpret directional cues [65, 66]. Coupling protrusion dynamics to external biochemical gradients and intracellular regulators, such as Cdc42, would enable future studies of how extracellular signals are integrated with intracellular biomechanical processes to shape collective cell migration patterns.

Lastly, future studies will be essential to better define properties of large cells and how they differ from and relate to regular-size cells. In our model, we made several specific assumptions about the biology of large cells. However, they will need to be characterized using comprehensive profiling methods, such as with single-cell RNA-sequencing and single-cell ATAC-sequencing to describe their transcriptome and genome-wide open *vs.* closed chromatin state, respectively. These sequencing methods are now commonly used and can be applied, in principle, to epithelial cell cultures featuring heterogeneous-size cells. Also, it will be important to perform an in-depth characterization of large *vs.* regular-size cell subcellular structure, including their organelle composition and abundance as well as their cytoskeletal make-up. The above additional molecular and supra-molecular data can then be input into the next generation of the model for more insightful and biologically-realistic predictive simulations on collective cell motion.

## 4 Methods

### 4.1 Model Description

To study dynamics of migrating cells, we employed the Subcellular Element (SCE) method, which has been widely applied in developmental biology [67, 36], plant stem cells studies [37, 68], and blood clotting dynamics [69] to name but a few.

In the SCE framework, each cell is represented by two sets of nodes, *N_m_* membrane nodes and *N_i_* internal nodes. In this study, we modeled two types of cells, large and regular-size cells that migrate and interact on a flat substrate. We used two-dimensional (2D) formulation to represent top-view projected morphology of cells. This formulation reduced computational cost relative to three-dimensional model and enabled direct comparison with experimental imaging data on collective epithelial cell migration on flat substrates [9, 12, 14], as well as with our own experimental imaging data on interactions between large and regular-size MDCK cells. This two-dimensional representation is also motivated by the main morphological distinction considered in this study that large cells occupy larger projected area on the substrate than regular-size cells. Yet, because large cells often exhibit flattened morphology, differences in projected area do not necessarily imply proportional differences in cytoplasmic volume [12]. Thus, the model isolates how area-associated differences in cell geometry, adhesion capacity, and motility can influence interactions between large and regular-size cells. The latter represent the “default” epithelial cell population calibrated using SCE framework, whereas large cells have increased projected area and node number. Unless otherwise specified, two cell types share the same baseline mechanical parameters, allowing differences in cell cluster behavior to be attributed to size-associated geometry, substrate coupling, and polarity-determination mechanisms. A schematic of the model is shown in Figure 9B.

**Figure 9:**
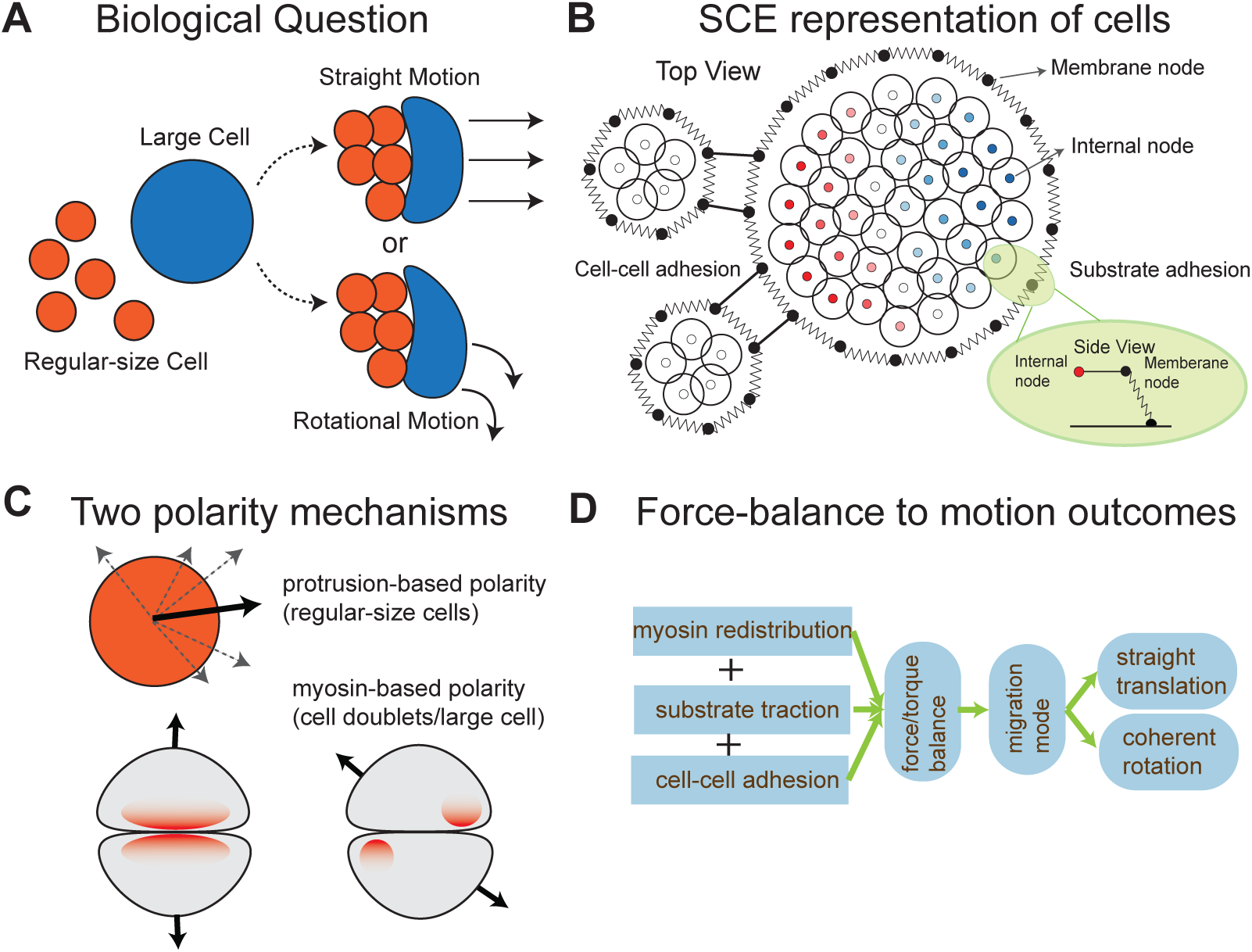
Description of the computational model and the biological question. (A) Diagram depicting biological setting and the question. Mixed-size epithelial cell cluster consists of one large cell and several regular-size cells. (B) Diagram of the Subcellular Element (SCE) model. It depicts one large cell and two regular-size cells. Points connected by springs represent membrane nodes, while nodes enclosed by circles represent internal nodes, with circles illustrating support of the Morse potential. Black solid lines between membrane nodes of different cells indicate cell-cell adhesion connections. Internal node colors in a large cell reflect myosin levels at the node, ranging from high (red) to low (blue). Light green inset shows side view of cell boundary and how cell-substrate adhesion sites are modeled. Note that this diagram is not obtained from numerical simulations and is for illustrative purposes only. (C) Two polarity mechanisms used in the model. Top: random protrusion extension defines regular-size cell polarity through a weighted directional sum. Bottom: contact-induced myosin redistribution reorients polarity away from the myosin-enriched contact region. (D) Flowchart summarizing model-predicted mechanism linking local cell-level interactions to cluster-scale group migration. Myosin asymmetry, substrate traction and cell-cell adhesion determine net force/torque balance, which in turn selects the migration mode between either straight translation or coherent rotation.

In this section, we provide detailed description of our 2D multi-scale model of cell motility and cell-cell interactions on a flat substrate based on the following hypothesized biological mechanisms:

[H1] For both large and regular-size cells, motility is driven by interactions with the substrate through dynamic assembly and disassembly of focal adhesions toward the polarity direction, defined below.
[H2] Each cell is assigned a polarity vector that determines its direction of movement. In the model with cellular protrusions, it is assumed that the weighted sum of protrusion directions determines cell polarity direction.
[H3] Cell polarity is also defined using the intracellular myosin distribution for systems consisting of adhered pairs of regular-size cells and for one large cell interacting with regular-size cells. In this case, upon attachment, myosin redistributes and accumulates near the cell-cell interface. We defined cell polarity as the outward normal at the membrane point farthest from the myosin peak. This assumption reflects well-established observation that myosin is typically concentrated toward the rear of migrating cells [28, 70, 71, 72]. Local myosin levels also influence disassembly rate of adhesions as defined below.

In the model, cell polarity was specified through two mechanisms. For isolated regular-size cells and for regular-size cells within clusters containing one large cell, polarity was governed by randomly generated protrusions that probe the surrounding environment. When no other cells were detected nearby, polarity vector was computed as the length-weighted sum of protrusion directions. If the center of another regular-size cell lay within one cell radius of the tip of a protrusion, that protrusion direction received additional weight in the polarity calculation. Similarly, when large cell was present, additional directional bias toward large cell was included. Another mechanism is the myosin-redistribution-dependent polarity specification as described in [H3].

In the following sections, we provide detailed description of each component of the model.

#### 4.1.1 Model components

##### Langevin equations describing dynamics of the SCE nodes

We followed similar approach as in the Epi-scale model [73, 67]. Nodes interact through a set of potential forces, springs and Morse potentials, that maintain volume and shape of each cell, as well as interactions between neighboring cells. Interactions between internal nodes in the same cell represent cytoplasmic pressure (*E*^II^), where pressure from cytoplasm onto membrane is modeled through interactions between internal nodes and membrane nodes (*E*^MI^). Interactions between membrane nodes within the same cell represent cortical stiffness (*E*^MMS^). Between two neighboring cells, adhesion is modeled through pairwise interactions between membrane nodes (*E*^adh^) and potential *E*^MMD^ is added to account for repulsive force between membrane nodes in different cells which prevents membranes from overlapping. Cell-substrate adhesions are modeled as springs with zero rest length as described later. Position of nodes evolves according to the following equations:

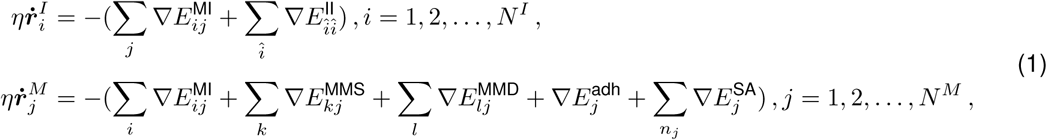

where *η* denotes damping coefficient, 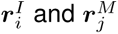 are positions of internal node *i* and membrane node *j*. In both equations, first summation runs over all membrane/internal nodes, while second summation in the equation of 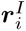 loops over all other internal nodes. *k* and *l* represent the index of membrane node, within the same cell or from a neighboring cell, interacting with the current membrane node *j*, respectively.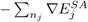 represents total force generated by cell-substrate adhesion connections attached to node *j* and indexed by *n_j_*. In the model, we impose that the number of adhesion connections per node is bounded above by 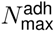. Explicit definition of this force term is provided below. Sub-model involving only node-node interactions was previously calibrated for epithelial cells in *Drosophila* wing disc [73, 67]. In the present study, same mechanical parameters are used for both large and regular-size cells, with the exception that bending coefficient of membrane nodes involved in the definition of *E*^MMS^ is slightly reduced, allowing large cell to occupy larger area and to be discretized with a greater number of nodes. Parameters related to cell polarity and motility are calibrated as described in the Parameter Calibrations section in SI. Note that equations in system (1) are not deterministic, as substrate-adhesion force generated by 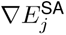 contains stochastic components arising from random assembly and disassembly of focal adhesions.

##### Cell polarity determination

Both regular-size and large cells are initialized as circular discs with radii *R*_r_ and *R*_l_, respectively. Cell polarity is represented by a unit vector ***p****_c_*(*t*), defined as ***p****_c_*(*t*) = (cos(*φ_c_*(*t*)), sin(*φ_c_*(*t*))), where *c* is the cell index and *φ_c_*(*t*) is the cell polarization angle measured counterclockwise from the positive *x* axis. Initial values of polarization angle are randomly selected from [0, 2*π*), and their time evolution is determined by intracellular myosin distributions or protrusion dynamics, together with positions of neighboring regular-size and large cells in clusters simulation, as described below.

##### Cell protrusion extension

During migration of single epithelial cells, actin-based protrusions, such as filopodia and lamellipodia, facilitate cell movement by probing environment and generating traction forces. In the context of epithelial cell monolayers, protrusions, such as lamellipodia, also contribute to maintenance of cell-cell contacts, playing critical role in coordinated collective migration [21, 74, 75, 76, 77]. Based on these observations, in the model we assume that regular-size cells can extend protrusions to probe environment and the direction of these protrusions influence direction of single cell motion [21, 78]. Each regular-size cell is able to randomly generate up to *N*_prot_ protrusions with the rate *P_p_*, maximum length *L*_max,p_, maximum lifetime *T_p_*, and orientation angle *φ_p_*, defined relative to the *x*−axis. After formation, protrusions extend and retract with their length governed by *L_p_*(*τ*) = *L*_max,p_(1 − cos(2*πτ/T_p_*)), where *τ* denotes age of the protrusion.

Direction of motion of each cell is regulated by the combined influence of its protrusions, as well as the spatial configuration of neighboring regular-size and large cells. If another regular-size cell lies near the tip of a protrusion, contribution of that protrusion to cell polarity is enhanced. This promotes aggregation of regular-size cells and formation of multi-cellular clusters. In the presence of a large cell, directional bias is introduced toward large cell. Taking together, polarization angle *φ_c_* of the *c*-th regular cell is governed by the following equation

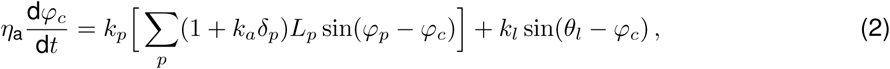

where *η_a_* is the damping coefficient. Parameter *k_p_* quantifies contribution of protrusion orientation to cell polarity, with each protrusion weighted by its length *L_p_*. *δ_p_* indicates whether the tip of a protrusion with orientation *φ_p_* lies near the membrane of a neighboring cell, and in this case contribution of that particular protrusion is amplified by factor *k_a_*. Parameter *k_l_* controls strength of repolarization toward large cell, whose position relative to current cell is given by angle *θ_l_*.

##### Myosin dynamics

In isolated cells, we assume that steady-state myosin distribution is uniform, representing a condition in which unbound myosin motors diffuse freely in the cytoplasm [79, 80]. Upon contact with another cell, activated myosin becomes enriched near the adhesion region, where it generates contractile forces essential for adherens junction reinforcement and stabilization [50]. In the present study, we model myosin transport among all internal nodes within each cell and incorporate myosin-dependent contractility into disassembly of adhesion bonds associated with each node. Myosin redistribution in the cytoplasm is described by introducing flux among internal nodes composed of two independent components, referred to as the symmetric and asymmetric components. Myosin dynamics at each internal node *i* are governed by weighted exchanges with neighboring nodes:

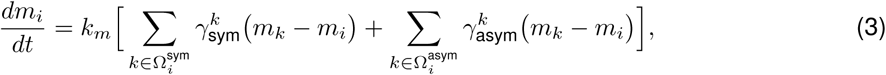

where *m_i_* denotes myosin level at node *i*, *k_m_* is myosin redistribution rate, and 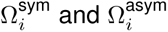 denote sets of neighboring nodes that can exchange myosin with node *i* in the symmetric and asymmetric cases, respectively. For simplicity, we assume that non-negative weights 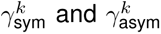 are uniform within each neighborhood, i.e. 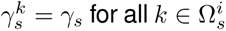 with *s* ∈ {sym, asym}. Symmetric and asymmetric components correspond to myosin fluxes that are uniformly directed toward cell-cell interface or biased toward one side of the interface, respectively. This decomposition reflects observed myosin recruitment and activation near cell-cell contacts and polarized myosin distribution [23].

For the symmetric component, 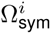 consists of nodes located within a disk of radius *d_s_* centered at node *i* that are farther from the adhered membrane region than node *i*. Node *i* has non-zero coefficients *γ_sym_* only if it lies within a distance *d*_tr,1_ of the adhered membrane region and its myosin level is below threshold *m*_max_. Distance *d_s_* is defined such that almost all internal nodes in a given cell, regular-size or large, are included. The asymmetric component, 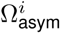, is defined similarly, except that distances are measured from a designated endpoint of the adhered membrane segment, identified as the last membrane node along interface in clockwise direction when viewed from cell interior. This definition introduces a directional bias in myosin flux along the interface. Such biased redistribution represents polarized contractile activity near cell-cell contact, which had been implicated in the rotation motion of small cell assemblies. Corresponding weights *γ*_sym_ and *γ*_asym_ are nonzero only for nodes belonging to their respective neighborhoods. We compute myosin level associated with each membrane node by averaging myosin values of internal nodes located within a distance *d*_tr,2_ of that membrane node. Cell polarity is then defined as the outward normal direction of the membrane at the location farthest from the peak of myosin computed at membrane nodes. In case of a cell doublet, when only symmetric component is active, myosin tends to spread uniformly along the contact interface, resulting in the absence of rotational motion. When only asymmetric component is active, myosin accumulates preferentially on one side of the interface, leading to sustained rotational motion. When both symmetric and asymmetric flux components are active, i.e. when *γ*_sym_ and *γ*_asym_ are both nonzero, symmetric term provides uniform enrichment of myosin along cell-cell interface, while asymmetric term introduces directional bias that drives preferential accumulation on one side of the contact. Consequently, myosin profile is partially polarized along the interface, with the degree of polarization controlled by the ratio *γ*_asym_*/γ*_sym_. Under mixed condition, rotational motion remains sustained but occurs at angular velocities intermediate between those in the purely symmetric and purely asymmetric cases. In case of mixed-size cell clusters, cell and cluster motion is significantly more complex, as shown in Section 2.

##### Focal adhesion assembly and disassembly

We model motility of large and regular-size cells toward the direction ***p****_c_* as a result of stochastic binding and unbinding of cell-substrate adhesions, following classical approaches on cell motility modeling [28, 29, 30]. It is assumed that force transmission between cell and substrate is primarily through the formation of integrin-based focal adhesions (FAs), which are mechanosensitive complexes that grow in response to increased mechanical tension, and diminish when tension is reduced. In the present study, we assume each membrane node with coordinate ***r*** = (*x, y*) can bind randomly to a point on the substrate. A candidate substrate point is generated as *x_s_* = *x* + *l_s_* cos(*β_s_*), *y_s_* = *y* + *l_s_* sin(*β_s_*), where *l_s_* is drawn from a truncated Gaussian distribution with mean *µ_l_* and standard derivation *σ_l_*, values of which are chosen so that the range of sizes of initial adhesion bond matches with the size of focal adhesions observed in experiments [81]. Angle *β_s_* is chosen within an angular range of ±*π/*4 around cell polarity direction. Probability that a specific adhesion forms during the time interval [*t, t* + *dt*) depends on the bond size *l_s_*, and is computed by 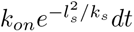 is the baseline formation rate and *k_s_* is scaling factor [82]. Each node can form up to 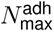 adhesion bonds, with the total number of allowable bonds for large cell and regular-size cell be *N_l_* and *N_r_*, respectively. An existing adhesion bond is modeled by a zero-length spring with stiffness *k*_SA_, connecting substrate point ***r****_s_* = (*x_s_, y_s_*) to node ***r*** = (*x, y*). A zero-length spring has zero equilibrium length and therefore exerts restoring force on the node ***r*** given by:

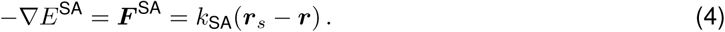

If spring becomes long or if local myosin level *m* at the node ***r*** is high, adhesion is likely to break. At every time step, an existing adhesion bond is removed with probability *p_b_* with *p_b_* = (1−*e*^−^*^k^*^off(^*^l/l^*^0 +^*^m/m^*^0)^*^dt^*), where *l* is the current spring length, and *l*_0_ and *m*_0_ are the characteristic values for spring extension and myosin level, respectively.

##### Computational implementation of the model

The model is implemented in C++ and parallelized using CUDA, with all simulations performed on graphics processing units (GPUs) from UC Riverside HPCC cluster. Mechanical forces that maintain cell integrity, regulate cell shape, and mediate cell-cell adhesion are updated following the algorithm described by Nematbakhsh et al. [73], with no cell division in the present study. Model components associated with polarity determination and cell-substrate adhesion are updated after evaluation of other mechanical forces and prior to updating node positions. Specifically, cell polarity is first determined based on either myosin distribution or protrusion dynamics, after which cell-substrate adhesion binding and unbinding are updated. Substrate-derived forces, together with other mechanical forces, then govern motion of each node. System of differential equations in Eq. (1) is solved numerically using standard explicit Euler scheme.

### 4.2 Model parameters

Model parameters and their values are shown in Table 1.

### 4.3 Experimental cell culture and time-lapse imaging

MDCK (NBL-2; CCL-34, ATCC) epithelial cells were cultured in Dulbecco’s modified Eagle medium supplemented with 10% fetal bovine serum and 1% penicillin-streptomycin. Cells were maintained at 37 ^◦^C in a humidified incubator containing 5% CO_2_. Culture medium was replaced three times per week, and cells were passaged upon approaching confluence.

For live-cell imaging, approximately 2 × 10^4^ cells were seeded in standard TC 12-well plate and allowed to attach and form spatially separated cells and small epithelial clusters. Multi-position time-lapse imaging was performed for up to 48 h using an Olympus FLUOVIEW FV3000 microscope. Cells were maintained at 37 ^◦^C and 5% CO_2_ throughout imaging. Images were acquired every 15 minutes.

Four representative time-lapse sequences capturing interactions between an enlarged MDCK cell and surrounding regular-size cells were selected for analysis and designated datasets A-D.

Enlarged cells were identified based on their greater projected area and flattened morphology relative to neighboring regular-size cells.

## Acknowledgments

Funding

MA was partially supported by the National Science Foundation grant DMS 2424826, MVP by the National Science Foundation grant IOS-2421118.

## **5** Data Availability Statement

The code used to run the simulations and generate the analyses is available at https://github.com/jiagou105/epithelial-cell-cluster.git.

## **6** Author Contributions

Conceptualization: Mark Alber, Maxim Plikus, Jia Gou, Mykhailo Potomkin

Model construction: Jia Gou, Mark Alber, Mykhailo Potomkin

Model calibration: Jia Gou, Mykhailo Potomkin

Simulation investigation: Jia Gou, Mykhailo Potomkin

Experimental investigation: Jonard P. Ingal, Sergei Butenko, Maxim Plikus, Wendy F. Liu

Image analysis: Mykhailo Potomkin, Jia Gou

Formal analysis: Jia Gou, Mykhailo Potomkin

Visualization: Jia Gou, Mykhailo Potomkin

Supervision: Mark Alber, Maxim Plikus, Sergei Butenko, Wendy F. Liu

Writing - original draft: Jia Gou, Mykhailo Potomkin, Mark Alber, Maxim Plikus

Writing - review & editing: Jia Gou, Mykhailo Potomkin, Mark Alber, Maxim Plikus, Sergei Butenko, Wendy F. Liu

## Supporting information captions

S1 Text. Analysis of isolated single-cell motion and calibration of model parameters.

**S1 Video. Representative dataset A of collective cell motion following contact with a large cell.** Phase-contrast time-lapse microscopy showing a group of regular-size cells contacting a large cell, followed by motion of the large cell and the resulting mixed-size cell cluster.

**S2 Video. Representative dataset B of collective cell motion following contact with a large cell.**

**S3 Video. Representative dataset C of collective cell motion following contact with a large cell.**

**S4 Video. Representative dataset D of collective cell motion following contact with a large cell.**

