## Supplemental Information for "Study of Principles Governing Epithelial Cell Clustering and Collective Motion *In Vitro*"

### 1 Supplemental Information

In the following sections, we describe the dependence of cell movement speed on cell-substrate interactions, and the analysis of single cell trajectories, as well as the parameter calibrations.

#### 1.1 Dynamic cell-substrate adhesion regulates epithelial cell motility by balancing traction generation and adhesion turnover

The cell-substrate adhesion and protrusion-based polarity modules are described in the main text. This coarse-grained adhesion representation is related to motor-clutch models, but here the combined cell-substrate mechanical coupling is represented by a single effective spring parameter $k_{\mathrm{SA}}$ [1, 2, 3]. Here, we isolated these modules in single-cell simulations to evaluate how adhesion dynamics and protrusion-driven polarity regulate autonomous cell motility. Specifically, we simulated an isolated regular-size cell and varied key adhesion parameters, including adhesion coefficient $k_{\mathrm{SA}}$, characteristic disassembly length $l_0$, baseline adhesion bond formation rate $k_{\mathrm{on}}$, and maximum number of adhesion sites per node $N_{\mathrm{max}}^{\mathrm{adh}}$. For each parameter set, all other parameters were held fixed at their baseline values, and mean cell speed was computed from the simulated cell trajectories.

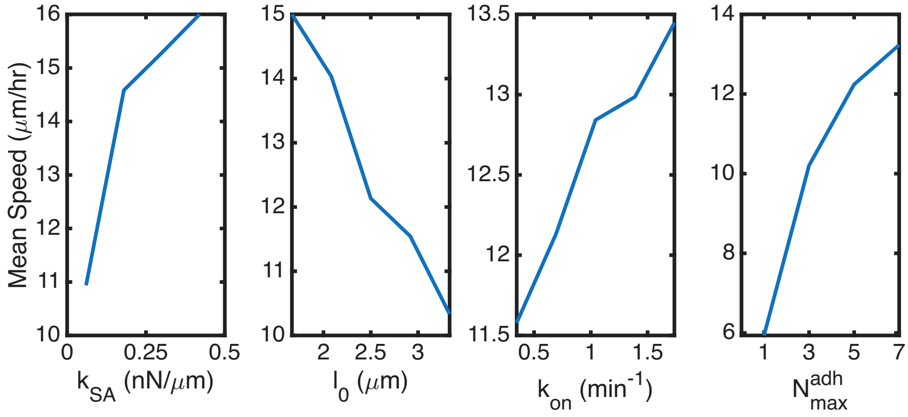


Fig. S1: Dependence of cell speed as functions of adhesion parameter $k_{\mathrm{SA}}$, characteristic unbinding length $l_0$, baseline adhesion formation rate $k_{on}$ and number of maximal adhesion sites $N_{\mathrm{max}}^{adh}$ can be formed at a single node.

As shown in Fig. S1, the mean cell speed shows positive relation with $k_{\mathrm{SA}}$, agreeing with the observations that cells are generally move faster on stiffer substrate [4, 5]. In addition, higher value of the characteristic unbinding length is associated with lower rates for focal adhesion disassociation and more stable bonds, which leads to lower migration speed. Cell speed also increases with increasing basal focal adhesion formation rate $k_{\mathrm{on}}$ and with the maximum number of adhesion sites, $N_{\mathrm{max}}^{\mathrm{adh}}$. Both parameters are directly related to the magnitude of the traction force transmitted at an individual node, thereby enhancing overall cell motility.

#### 1.2 An isolated regular-size cell exhibits persistent random-walk behavior

Experimental studies have shown that epithelial cell migration follows anomalous diffusion dynamics, instead of normal Brownian motion [6], as cell motility is driven by active biological processes rather than random thermal fluctuations. Within our modeling framework, cell polarity is determined by the emergence and turnover of cellular protrusions, and migratory forces are generated through assembly and disassembly of adhesion bonds. We then study whether these two components are sufficient to reproduce the anomalous diffusion observed in the motion of isolated cells.

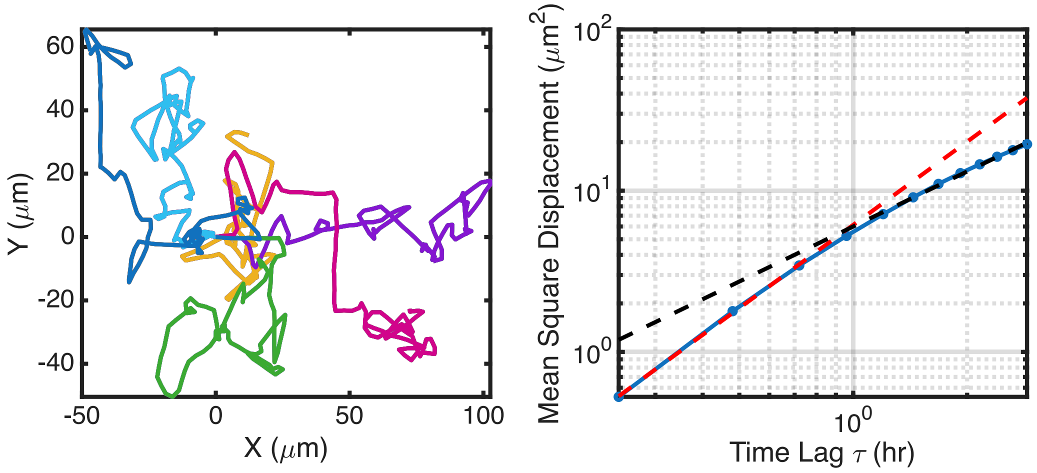


Fig. S2: (A) Migratory trajectories of single isolated regular-size cells (5 independent simulations). (B) Mean squared displacement (MSD) of one single isolated regular-size cell with the parameters used in (A). The fitted red and black dashed lines have slope of 1.71 and 1.13, respectively.

Fig. S2a shows typical migratory trajectories of a single isolated cell. For one isolated cell, the mean squared displacement (MSD) is computed using the following expression

$$MSD(\tau) = \frac{1}{N-\tau} \sum_{i=1}^{N-\tau} \left( (x_{i+\tau} - x_i)^2 + (y_{i+\tau} - y_i)^2 \right),$$

where $(x_i, y_i)$ is the cell location at time step $i$, $N$ is the total number of time steps and $\tau$ is the time lag. The average MSD is then computed as the mean of MSDs obtained from individual trajectory and the resulting curve is shown in blue curve on the right panel of Fig. S2. To characterize the diffusion, we also fit the average MSD with the following curve

$$MSD(\tau) = K\tau^{\alpha}.$$

The fitted line, for small lag, exhibited a slope $\alpha > 1$, which is consistent with the observation that cell motion is better described by superdiffusion rather than classical Brownian motion. Additionally, for time lags shorter than an hour, the fitted value of $\alpha$ is $\sim 1.71$ which decreases to $\sim 1.13$ as $\tau$ increases. This behavior is consistent with previous studies suggesting that a "persistent random walk model" more accurately captures the average MSD of migrating cells as a function of time lag $\tau$, and the fitted parameter $\alpha$ agrees with that obtained from fitting experimental data [7, 6]. In the persistent random walk model, MSD first exhibits ballistic behavior characterized by $\alpha \approx 2$ for small lag $\tau$ and transitions to $\alpha \approx 1$ for large lag. In our simulation of single isolated cells, only two mechanisms are included: the stochastic extension of protrusions, which determines migratory direction, and focal adhesion assembly and disassembly mediating cell-substrate bindings. The above analysis showed that this simple model is sufficient to reproduce the superdiffusive behavior experimentally observed in single-cell migration at small lag times and normal diffusion at larger time lags. In addition, this model provides a framework for incorporating additional subcellular features and regulatory mechanisms that influence cell motility.

Next we investigate how the alignment rate $k_p$ of cell angle with net protrusion direction, protrusion extension rate $P_p$, protrusion lifetime $T_p$ affect the migratory behavior of a single cell. In Fig. S3a, the blue curve denotes dependence of the fitted $\alpha$ at small lag times on the corresponding parameter, while the red curve denotes that of the fitted $\alpha$ at large lag times. The fitted MSD exponents indicated persistent, superdiffusive motion at short lag times and a crossover toward nearly diffusive behavior at long lag times. Among the three parameters, the protrusion extension rate $P_p$ has the strongest influence: increasing $P_p$ substantially decreases both short- and long-time exponents, indicating reduced polarity persistence and faster decorrelation of motion. In contrast, the alignment rate $k_p$ primarily modulates the long-time exponent, with stronger alignment rate reducing long-time persistence while only weakly affecting short-time behavior. The protrusion lifetime $T_p$ has a comparatively modest effect, mainly providing a fine adjustment of long-term memory with little change in the short-time exponent. Together, these results show that the model consistently exhibits short-time ballistic and long-time diffusive motions. The MSD scaling is controlled by protrusion-driven polarity persistence, with $P_p$ acting as the dominant parameter and $k_p$ and $T_p$ playing secondary roles.

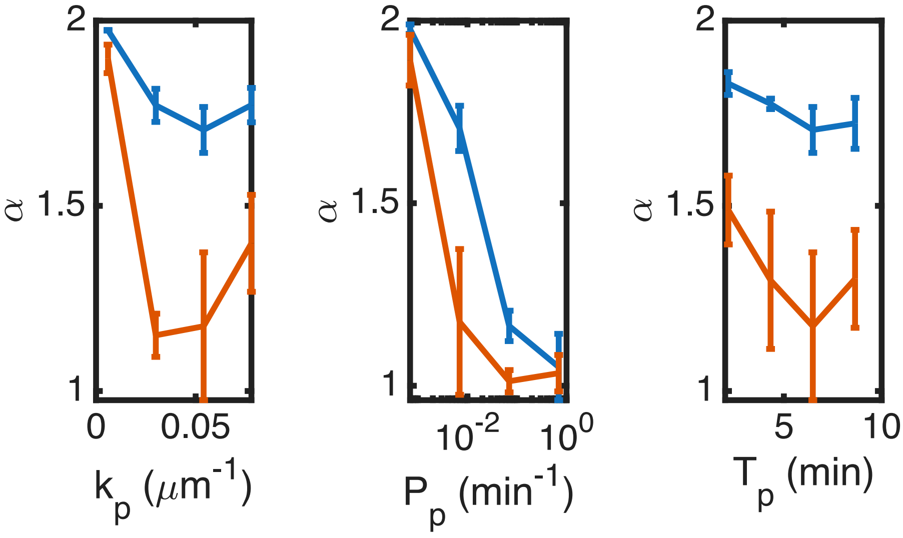


Fig. S3: The fitted values of $\alpha$ for small (blue) and large (red) lag times $\tau$, shown as functions of the alignment rate of cell angle with net protrusion direction $k_p$, protrusion extension rate $P_p$, and lifetime of cellular protrusions $T_p$.

Overall, the above results showed that single-cell migration persistence emerges from the competition between protrusive polarity memory and adhesion turnover.

#### 1.3 Parameter Calibrations

The model parameters determining the node-node interactions, including the Morse potential parameters in the energy functions $E^{\mathrm{II}}$, $E^{\mathrm{MI}}$ and $E^{\mathrm{MM}}$, follow values used for similar epithelial cell in prior modeling of the *Drosophila* wing disc [8]. In this section, we discuss choices of model parameters related to cell motility.

Large cells can spontaneously emerge within epithelial cells colonies on a flat substrate and have been reported to exhibit substantially larger spread areas than regular-size cells. Epithelial cells typically have radii in the range of $\sim 5-18$ $\mu m$ [9, 10]. Here we take a representative radius $R_{\mathrm{r}} = 15$ $\mu m$ for regular-size cells and choose $R_{\mathrm{l}} = 50$ $\mu m$ to represent a highly spread large cell.

Cell polarity determination is modeled using two mechanisms: protrusion-mediated directional sensing, and contact-induced myosin redistribution. For protrusion-mediated polarity, each regular-size cells is allowed to carry at most $N_{\mathrm{prot}} = 5$ simultaneous protrusions. Quantitative imaging studies of other cell types indicated that the instantaneous number of filopodia per cell typically lies in a range of several to tens, and the value chosen here is to capture the localized exploratory activity in a phenomenological manner, rather than an exact count of protrusions [11]. Reported filopodia and lamellipodial extension lengths in epithelial cells are typically on the order of $1-16$ $\mu m$. We therefore set the maximum protrusion length to $L_{\mathrm{max,p}} = 5$ $\mu m$ to represent a length scale that facilitates protrusive sensing and promotes cell-cell encounters [12, 13]. Experimental studies in keratinocytes and other mammalian cells have reported filopodial lifetimes ranging from seconds to several minutes [14, 15, 16]. Here we choose a protrusion lifetime $T_p = 2-10$min which lies within the experimentally observed range and allows sufficient time for protrusions to probe the local environment [17].

In addition to protrusion-mediated mechanism, another mechanism is that cell polarity is influenced by contact-induced myosin redistribution toward cell-cell interface, which was incorporated in the cell doublets and mixed cluster models. As quantitative measurements of intracellular myosin levels are not available, myosin in the model is treated as an effective activity variable, and this approach focuses on relative spatial redistribution rather than absolute levels. We define a threshold distance $d_{\mathrm{tr},1} = 13.3$ $\mu m$ between an internal node and the cell-cell contact curve required to generate myosin flux toward that node. This distance is smaller than the regular-size cell radius and restricts contact-induced myosin recruitment to regions proximal to the contact interface. This is consistent with experimental observations that myosin accumulation and contact-associated contractile activity are localized near cell-cell junctions and their immediate vicinity.

Cell-substrate interactions are described by the dynamic assembly and disassembly of adhesion bonds, which are represented as elastic springs. In epithelial cells, focal adhesion clusters have been measured to have typical lengths of $2.05 \pm 0.57$ $\mu m$ [18]. In our model, the initial extensions of adhesion springs are drawn from a truncated Gaussian distribution with mean $\mu_{\mathrm{adh}} = 1.845$ $\mu m$ and standard deviation $\sigma_{\mathrm{adh}} = 0.513$ $\mu m$, chosen so that the majority of bond lengths fall within the range of 1.48 $\mu m$ to 2.62 $\mu m$, in accordance to experimental observations.

The force transmitted between a cell and the substrate is influenced not only by focal adhesion size but also by the molecular composition of focal adhesions, associated signaling pathways, and substrate properties, including stiffness. In the present model, these factors are captured using a single effective parameter $k_{\mathrm{SA}}$, which represents the stiffness of the linear spring describing the cell-substrate linkage. It provides a coarse-grained representation of the molecular bonds that connects cell and substrate. The range of $k_{\mathrm{SA}}$ was chosen to be $0.006-0.42$ $nN/\mu m$. For adhesion extensions on the order $1.48-2.62$ $\mu m$, this corresponds to forces of approximately $\sim 10^{-2}-1$ $nN$ per adhesion bond. Assuming that at most $N_{\mathrm{max}}^{\mathrm{adh}} = 10$ adhesion bounds per node and approximately 80 membrane nodes for a regular-size cell (or $\sim 240$ for a large cell), the resulting total traction force can reach the tens-to-hundreds of nN range, depending on the fraction of engaged adhesions and their extensions. This range is broadly consistent with experimentally measured traction forces in epithelial cells [19]. The baseline adhesion binding rate $k_{on}$ and the adhesion area scale $k_s$ together determines the effective binding rate of individual adhesion sites. Adhesion formation is evaluated at every simulation update corresponding to a small physical time step ($dt = 0.026$ sec) and these parameters are chosen to represent a microscropic attempt rate, while ensuring the per-update formation probability remains below one. The adhesion rupture rate is modeled as depending on both the adhesion spring length and the local myosin level, characterized by a length scale $l_0 = 1.67 - 3.33\ \mu m$ and a myosin scale $m_0 = 0.5$, respectively, together with an additional rate constant $k_{\text{off}}$. Myosin is treated as a non-dimensional variable with an initial basal level $m_{\text{init}} = 0.5$ at internal nodes and a typical maximal recruitment of $m_{\text{max}} = 3 - 6$ near the cell-cell contact. We choose $m_0 = 0.5$, equal to the basal myosin level, so that the myosin term in the exponent is of order one at rest ($m/m_0 \approx 1$) and increases rapidly as local myosin accumulates ($m/m_0 \approx 6 - 12$ near contact interface). This choice provides a coupling between actomyosin contractility and adhesion turnover as observed in experiments, and ensures that adhesions in low-contractility regions remain relatively stable and those under increased myosin activity are more likely to fails.
